# Mitotic bookmarking by Prox1 preserves mammalian neuronal lineage identity memory via promoting timely H3K27me3 restoration

**DOI:** 10.64898/2026.02.25.707603

**Authors:** Chouin Wong, Jie Liu, Haoran Yang, Haotian Li, Xiaoqi Luo, Tingyi Li, Zili Chen, Jingyi Chu, Yuying Shen, Shuai Long, Yong Zhang, Yan Song

## Abstract

Mitosis poses a daunting challenge to transgenerational inheritance of cell identity memory. How neuronal lineage identity is faithfully transmitted across mitosis remains largely unexplored. Here we report that, in mouse hippocampus, the transcription factor Prox1 acts as a mitotic bookmark for safeguarding neuronal lineage identity across cell divisions. Prox1 exhibits mitotic retention in dentate gyrus (DG) neural stem cells and defines DG lineage identity by suppressing the alternative cornu ammonis (CA) identity. Mitotic bookmarking by Prox1 at key bivalent CA identity genes promotes timely and precise restoration of Polycomb repressive complex 2 (PRC2)-mediated H3K27me3 deposition to avoid ectopic expression of CA identity genes and lineage identity crisis. Remarkably, specific mitotic retention-deficient *Prox1* conditional knock-in mice produce severe DG developmental defects. Thus, mitotic bookmarking imprints neuronal lineage identity in mouse hippocampus, which is likely to represent a fundamental principle underlying the preservation of lineage identity memory in mammalian brain development.

## Introduction

The astonishing complicity of our brain is built upon the mitotic divisions of a limited number of neural stem cells (NSCs), so called radial glial cells (RGCs), and neural progenitors that generate neurons and non-neuronal cells of enormous number and diversity^1^. In most of the cases, a specific neuronal lineage identity is initially defined in NSCs or progenitors by their spatial location and their expression of distinct temporal factors^2–5^. Such lineage identity is transmitted to their progenies through mitotic divisions to produce distinct neuronal subtypes of characteristic morphology and function.

Mitosis, however, poses a daunting challenge to the transgenerational inheritance of cell identity. In mitosis, when chromatin is highly condensed into chromosomes, the gene regulatory machinery is largely dispelled from chromosomes or degraded^6^. As a consequence, transcription is essentially halted^7–9^. It remains obscure how specific gene regulatory programs defining unique lineage identities are faithfully transmitted from neural stem cells to their progenies across cell divisions and then solidified in post-mitotic neurons.

Transcription factors and chromatin remodelers that are retained on mitotic chromosomes have been posited as mitotic bookmarks^8,10–14^. In mammalian cultured cells, these mitotic bookmarking factors have been found to specifically label key target genes and foster their timely transcriptional reactivation, ensuring efficient and precise reconstitution of specific gene regulatory programs upon mitotic exit and thereby transmission of cell fate memory across mitosis. Latest studies revealed that mitotic bookmarking is crucial for preserving neural stem cell fate memory in fly neural development^15,16^. However, whether mitotic bookmarking plays an important role in preserving neuronal lineage identities in mammalian brain development remains unexplored.

The hippocampus is a brain region crucial for social memory and spatial navigation^17–19^. The adult hippocampus is mainly composed of two interlocking parts: the dentate gyrus (DG) region primarily composed of the DG granule neurons and the cornu ammonis (CA) region mainly consisting of the CA pyramidal neurons^20^. During hippocampal neurogenesis, CA NSCs residing in the hippocampal neuroepithelium undergo asymmetric divisions to produce CA progenitors, which in turn differentiate into CA neurons^21^. In comparison, DG NSCs asymmetrically divide to produce DG progenitors, which migrate along the dentate migratory stream (DMS) until reaching the hilus, where they differentiate into DG neurons^4,22,23^. It remains enigmatic how DG and CA NSCs establish and maintain their distinct neuronal lineage identities.

The evolutionarily conserved homeobox transcription factor Prospero Homeobox 1 (Prox1) is highly expressed in mammalian DG granule neurons and plays an essential role in DG neuron formation and maintenance^4,20^. *Prox1* gene transcription in DG is induced by the Wnt morphogen, which forms a descending signaling gradient from DG to CA neuroepithelium^4,24–26^. However, at molecular level, how Prox1 regulates DG neuronal lineage identity establishment and maintenance remains largely unresolved.

The Polycomb repressive complex 2 (PRC2) promotes local chromatin compaction and epigenetic silencing of its target genes by adding methyl groups to histone H3 at lysine 27 (H3K27me3)^27–33^. The core subunits of PRC2 include enhance of Zest homolog 2 (EZH2), suppressor of Zeste 12 (SUZ12), embryonic ectoderm development (EED) and RB-binding protein 4/7 (RBBP4/7)^34–36^. Whereas the chromatin remodeling complex SWI/SNF (SWItch/Sucrose Non-Fermentable) has been found to act as mitotic bookmarks in mammalian cultured cells^11^, whether other chromatin remodelers such as PRC2 play a role in preserving cell fate or identity memory via mitotic bookmarking remains obscure.

Here we report that, in mouse hippocampal development, Prox1 acts as a mitotic bookmark for preserving hippocampal neuronal lineage identity memory across cell divisions. We find that Prox1 exhibits mitotic retention in DG but not CA NSCs and defines DG lineage identity by suppressing the alternative CA identity. Through establishing and performing technically-challenging *in vivo* low-input mitotic CUT&Tag with purified mouse DG NSCs, we reveal that Prox1 mitotically bookmarks key bivalent CA identity gene *Fezf2* to ensure timely and precise restoration of PRC2-mediated H3K27me3 deposition and thereby avoid ectopic expression of CA identity genes and lineage identity crisis. Remarkably, specific mitotic retention-deficient Prox1 conditional knock-in mice produce severe DG developmental defects, pinpointing the functional significance of Prox1 mitotic bookmarking. Notably, to our best knowledge, Prox1 is the first *bona fide* mitotic bookmark identified in developing mammalian brains. Our results highlight the pivotal role of mitotic bookmarking in preserving neuronal lineage identity memory across cell generations, providing new insight into neurodevelopmental disorder therapies^1,37^.

## Results

### NSC lineage-specific knockout of *Prox1* leads to severe DG malformation and spatial memory deficits

To investigate how the hippocampal neuronal lineage identity is specified and transmitted through cell divisions, we fist assessed whether Prox1 plays a pivotal role in establishing and maintaining DG lineage identity. We induced NSC lineage-specific *Prox1* knockout (KO) with tamoxifen-inducible *Nestin-CreER^T2^*. Tamoxifen was injected at E13.5 (embryonic day 13.5) and the *Nestin-CreER^T2^*; *Prox1^fl/fl^* (referred to as *Prox1^Nes-cKO^*) animals were examined at P0 (postnatal day 0) and P120 (postnatal day 120). Compared with the *Prox1^fl/fl^* littermate control, the DG region of *Prox1^Nes-cKO^* mice at P0 barely expresses Prox1 or DG immature neuronal marker Calb2 (Calretinin) (Supplementary Fig. 1a–g)^38,39^, indicating defective specification of DG lineage identity upon *Prox1* knockout. Not surprisingly, the DG but not CA region in *Prox1^Nes-cKO^* mice at P120 shrinks remarkably to around 25% of its normal sizes (Fig. 1a, Supplementary Fig. 1h, i). More importantly, in comparison to the littermate control, *Prox1^Nes-cKO^*mice at P120 exhibit drastic spatial learning and memory deficits (Fig. 1b, Supplementary Fig. 1j–l). Together, Prox1 is crucial for DG lineage identity specification and maintenance.

**Fig. 1.**
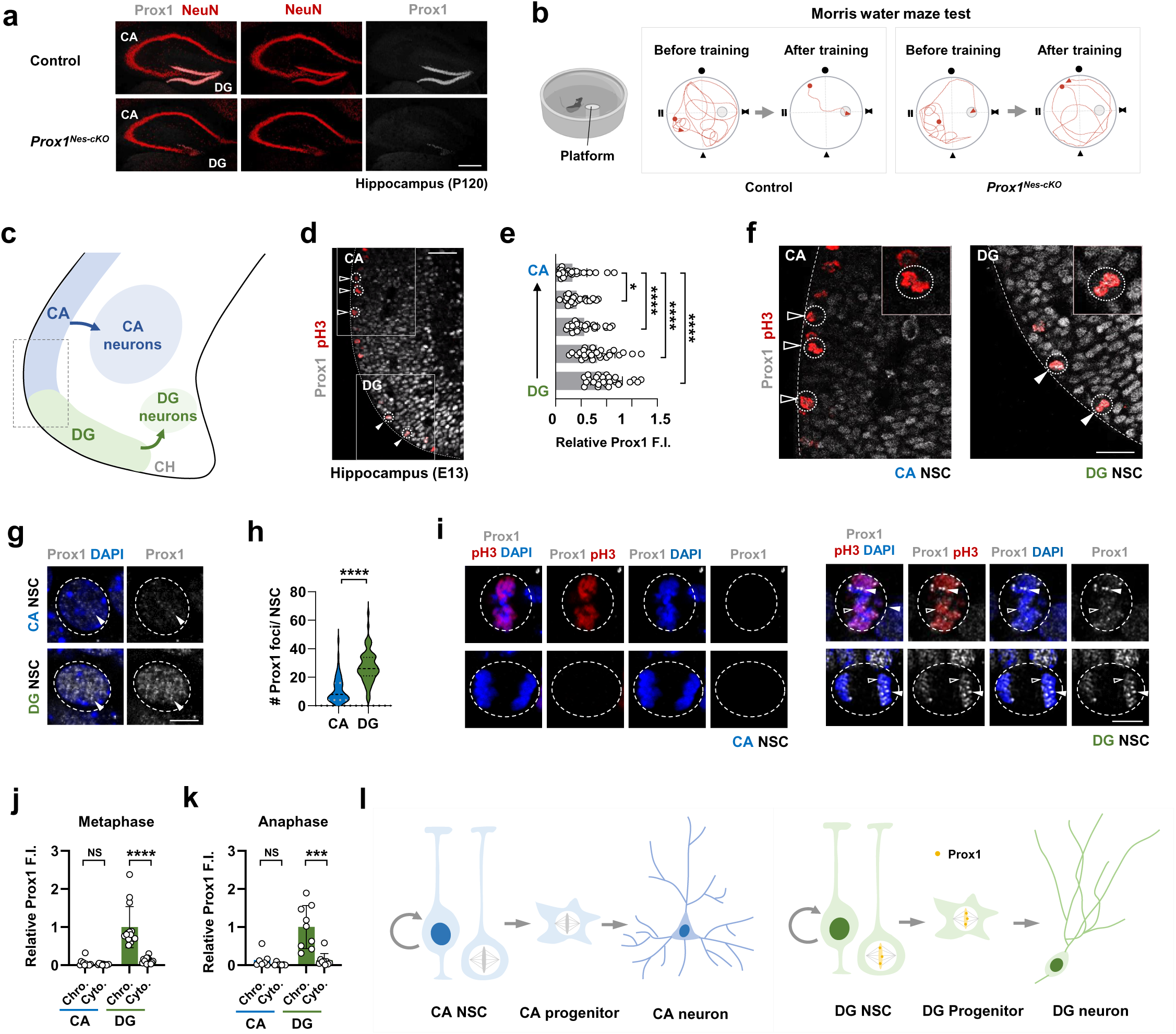
Prox1 exhibits mitotic retention in DG but not CA NSCs in embryonic hippocampus. (**a**) Representative images of control (*Prox1^fl/fl^*) and *Prox1^Nes-cKO^* (*Nestin-CreER^T2^*; *Prox1^fl/fl^*) hippocampus at P120 stained for Prox1 and NeuN (Neuronal nuclei), a pan-neuronal marker. Scale bar, 200 μm. (**b**) Representative swimming paths of the control and *Prox1^Nes-cKO^* mice before and after training in Morris Water Maze test. (**c**) A schematic diagram showing CA and DG neuroepithelium near the ventricular surface, with the neurogenesis process (arrows) producing CA and DG neurons respectively. (**d–f**) Representative images of developing hippocampus at E13 stained for Prox1 and pH3 (**d**, **f**) and quantifications of the relative Prox1 expression levels from CA to DG (**e**; n =27 brain sections). High magnification of the boxed areas in (**d**) showing the distribution of Prox1 in dividing CA or DG NSCs (**f**). *p < 0.05, ****p < 0.0001; Student’s t-test. Scale bar, 50 μm (**d**); 25 μm (**f**). The cell body of CA and DG NSCs is encircled by dotted lines. (**g**, **h**) Sample images of CA and DG NSCs in interphase stained with anti-Prox1 antibody and DAPI (**g**) and quantifications of Prox1 foci per NSC (**h**). n = 47, 60 cells; ****p < 0.0001; Student’s t-test. Scale bar, 5 μm. In this and subsequent micrographs, the cell body of CA and DG NSCs is encircled by dashed lines. (**i–k**) Representative images of metaphase (top) and anaphase (bottom) CA (left) and DG (right) NSCs at E13 stained with anti-Prox1, anti-pH3 and DAPI (**i**) and quantifications of Prox1 fluorescent intensity (F.I.) on the chromosomes (chro.) or in the cytosol (cyto.) (**j**, **k**). Arrowheads indicate Prox1 associating with chromosomes. n = 7, 9, 11, 14 cells, respectively; NS, not significant, ****p < 0.0001, ***p < 0.001; Student’s t-test. Scale bar, 5 μm. (**l**) A schematic diagram showing Prox1 (orange) mitotic retention in DG but not CA NSCs and progenitors.

### Mitotic retention of Prox1 in DG NSCs in embryonic hippocampus

We next examined the distribution pattern of Prox1 in the hippocampal primordium. Unexpectedly, we found that Prox1 is weakly but specifically expressed in the hippocampal epithelium as early as E13 and its distribution displays a descending gradient from DG to CA neuroepithelium (Fig. 1c–f). Prox1 forms discrete nuclear condensates in DG NSCs (Fig. 1g, h). In comparison, the expression levels of Prox1 in CA NSCs are significantly weaker (Fig. 1g, h). Intriguingly, Prox1 protein is retained on the mitotic chromosomes of the DG NSCs at E13, with spherical condensates enriched at pH3^-^ pericentromeric heterochromatin regions (Fig. 1f, i–k). In contrast, although Prox1 is weakly expressed in the CA NSCs, it fails to retain on their chromosomes (Fig. 1f, i–k). Furthermore, Prox1 is mitotically retain in dividing DG progenitors at E15.5, immediately before these progenitors differentiating into DG neurons (Supplementary Fig. 1m–o). Taken together, Prox1 is mitotically retained in DG NSCs and progenitors (Fig. 1l).

### DG and CA NSCs express distinct lineage identity genes in early hippocampus

To assess the functional significance of Prox1 mitotic retention in DG NSCs, we first sought to perform transcriptome analysis of DG vs. CA NSCs/progenitors at E13.5 and E14.5. The RNA-seq analysis revealed a series of <u>d</u>ifferentially <u>e</u>xpressed <u>g</u>enes (DEGs) in DG vs. CA NSCs/progenitors (Supplementary Fig. 2a). Some of the DG or CA DEGs are known to play crucial roles in hippocampal development or homeostasis. For example, Forkhead Box G1 (FOXG1) has been found to act in the CA lineage to repress DG neuronal identity^39,40^, whereas NeuroD6 is a well-characterized CA neuronal identity marker^41^. On the other hand, CXC Motif Chemokine Receptor 4 (CXCR4) is a chemokine receptor essential for DG development and maintenance^42^ and Hopx a unique homeodomain protein exclusively expressed in DG NSCs and progenitors^22,43^.

We next investigated how the transcription of these DEGs is altered upon NSC lineage-specific knockout of *Prox1*. We first used *Nestin-CreER^T2^* to induce NSC lineage-specific *Prox1* knockout in early hippocampal development. Following tamoxifen administration, we dissociated and isolated GFP^+^ *Prox1^Nes-cKO^* cells via fluorescence-activated cell sorting (FACS) at E18.5 and performed bulk RNA-seq (RNA sequencing). Interestingly, *Prox1^Nes-cKO^* cells not only downregulate DG-signature genes such as *Calb2* and *NeuroD4*^44^, but also upregulate CA-signature genes such as *FOXG1* and *Ctip2* (*Bcl11b*) (Supplementary Fig. 2b)^45^. In addition, the transcriptome profile of the *Prox1^Nes-cKO^* DG cells resembles that of CA lineage cells at E18.5 (Supplementary Fig. 2b), indicating a DG-to-CA lineage identity transition upon *Prox1* depletion.

We then induced *Prox1* knockout at an even earlier stage using *Emx1-Cre*, which initiates its expression in hippocampal NSCs starting E9.5^46^. We dissociated and isolated E15.5 control versus *Emx1-Cre*; *Prox1^fl/fl^* (referred to as *Prox1^Emx1-cKO^*) at DG cells and performed RNA-seq. Intriguingly, a different set of CA DEGs, including FEZ Family Zinc Finger 2 (*Fezf2*)^47–49^, is upregulated in *Prox1^Emx1-cKO^* DG cells (Supplementary Fig. 2c). This suggests that, upon *Prox1* depletion, there are at least two temporal waves of CA-signature genes that are upregulated sequentially. In accordance, a different set of DG DEGs, including *Hopx*, is downregulated in *Prox1^Emx1-cKO^*DG cells. Together, Prox1 defines DG lineage identity by simultaneously promoting DG identity and repressing the alternative CA identity.

We next explored whether Prox1 is sufficient to confer DG lineage identity. GFP-Prox1 is ectopically expressed in CA NSCs at E13 via *in utero* electroporation (IUE). At E14.5, cells expressing intermediate levels of GFP-Prox1 are enriched for transcriptome analysis. In comparison with control cells that express GFP, ectopic expression of GFP-Prox1 in CA NSC lineages is sufficient to induce ectopic expression of DG DEGs such as *Wnt5a* and substantially inhibit transcription of CA-signature genes such as *NeuroD6* and *Fezf2* (Supplementary Fig. 2d). Together, our results indicated that Prox1 is both necessary and sufficient to confer DG neuronal lineage identity by simultaneously inhibiting CA identity and promoting DG identity. CA-DEGs upregulated in *Prox1^cKO^*DG cells and DG-DEGs downregulated in *Prox1^cKO^* DG cells are therefore designated as CA identity genes and DG identity genes respectively (Supplementary Fig. 2e).

To further investigate how the DG vs. CA neuronal lineage identities are established, we performed single-cell RNA sequencing (scRNA-seq) on microdissected hippocampus from wild type E15.5 mouse embryos. Unsupervised clustering of high-dimensional transcriptome revealed 10 transcriptionally distinct cell clusters, including NSCs, progenitors, DG and CA neurons (Supplementary Fig. 2f). RNA velocity analysis and violin plot further confirmed that DG and CA lineages preferentially express DG and CA identity genes respectively (Supplementary Fig. 2g–i).

### Prox1 is mitotically retained at the key CA identity gene *Fezf2*

To identify Prox1 target genes in DG NSCs, we microdissected E13.5 dentate primordial regions and performed *in vivo* CUT&Tag assay. We noted that Prox1 preferentially binds to CA identity genes, such as *NeuroD6* and *Fezf2*, as well as DG identity genes, such as *Hopx* and *Prox1* itself (Fig. 2a, b). Collectively, our results indicated that Prox1 defines the DG lineage identity by binding and inhibiting the transcription of alternative CA lineage identity genes.

**Fig. 2.**
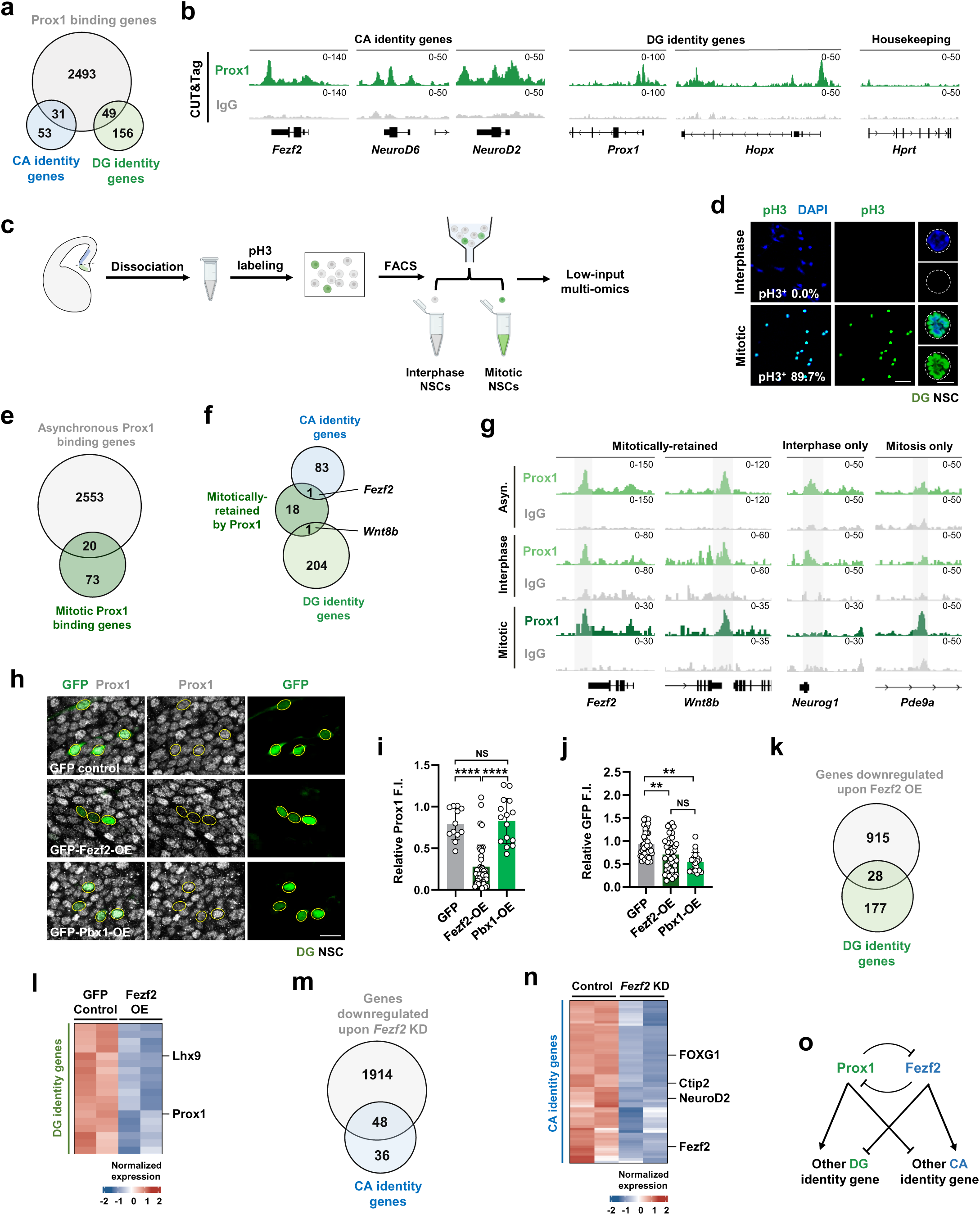
Prox1 is mitotically retained at the key CA identity gene *Fezf2*. (**a**) Venn diagram showing the intersections of Prox1 binding and DG and CA identity genes. (**b**) Genome browser examples of CUT&Tag for Prox1 tracks at CA and DG identity gene loci and housekeeping gene locus in DG NSCs at E13.5. (**c**) Experimental workflow of purification of mitotic and interphase NSCs from developing hippocampus followed by low-input multi-omics analysis. (**d**) Purity of the interphase and mitotic DG NSCs before and after FACS purification. pH3 indicate mitotic NSCs. n=99 (interphase) and 144 (mitotic), respectively. Scale bar, 100 μm (left); 5 μm (right). (**e**) Venn diagrams showing the overlap of Prox1 binding in asynchronous and mitotic DG NSCs. (**f**) Venn diagrams showing the intersection of Prox1 mitotic retention sites and CA or DG identity genes. (**g**) Genome browser examples of CUT&Tag for Prox1 and IgG tracks at *Fezf2*, *Wnt8b*, *Neurog1* and *Pde9a* gene locus in asynchronous (asyn.), interphase and mitotic DG NSCs. (**h–j**) Representative images of DG NSCs at E14.5 overexpressing GFP (top, control), GFP-Fezf2 (middle) or GFP-Pbx1 (bottom) stained for Prox1 (**h**) and quantifications of relative Prox1 (**i**; n=12, 41, 15 cells) and GFP (**j**; n=47, 43, 20 cells) expression levels. ****p < 0.0001, **p < 0.01; NS, not significant. Student’s t-test. Scale bar, 10 μm. (**k**, **l**) Venn diagram (**k**) and heat map (**l**) showing DG identity genes downregulated upon Fezf2 overexpression in DG NSCs. (**m**, **n**) Venn diagram (**m**) and heat map (**n**) showing CA identity genes downregulated upon *Fezf2* knockdown (KD) in CA NSCs. Scramble shRNA serves here as negative control. (**o**) Mutual repression between Prox1 and Fezf2, master DG and CA identity gene respectively.

In order to capture the genomic loci mitotically-bound by Prox1 in dividing DG NSCs, we developed a new pipeline suitable for enriching mitotic and interphase NSCs from developing mouse brains, allowing the identification of mitotic binding sites under physiological conditions (Fig. 2c). Briefly, mouse embryonic DG regions at E13.5 is microdissected and dissociated into single-cell suspensions, mildly fixed and immunolabeled with phospho-histone H3 (pH3) antibody before being sorted to enrich interphase versus mitotic DG NSCs (Fig. 2c). Our new pipeline allows interphase and mitotic DG NSCs with high purity (100% purity for interphase DG NSCs and 89.7% purity for mitotic NSCs) to be acquired before low-input multi-omics analysis being carried out (Fig. 2d).

After performing *in vivo* anti-Prox1 low-input CUT&Tag, we identified the binding sites of endogenous Prox1 in interphase and mitotic DG NSCs respectively. Around 0.77% Prox1 interphase binding sites are retained by Prox1 in mitotic DG NSCs (Fig. 2e, f), indicating that only a limited number of Prox1 target genes are selectively bound by Prox1 in mitosis. Remarkably, the CA identity gene *Fezf2* and the homeobox transcription factor *Pbx1* (*Pre-B-cell leukemia transcription factor 1*) are mitotically bound by Prox1 in DG NSCs (Fig. 2g).

### Fezf2 specifies CA lineage identity by suppressing the alternative DG identity

To understand why *Fezf2* but no other CA identity genes are selectively bound by Prox1 in mitotic DG NSCs, we first attempted to examine whether overexpression of Fezf2 might suppress *Prox1* gene expression. Pbx1, which is mitotically bound by Prox1 but not a CA identity factor, serves as a control. GFP-Fezf2, GFP-Pbx1 or GFP control is ectopically expressed in DG NSCs via IUE. At E14.5, immunofluorescence results indicate that, in comparison with control cells, ectopic expression of Fezf2 but not Pbx1 in DG NSC lineages leads to marked suppression of Prox1 levels (Fig. 2h–j), hinting that Fezf2 specifies CA lineage identity by inhibiting DG identity. To further test this notion, cells expressing low levels of GFP-Fezf2 at DG are enriched for transcriptome analysis. In comparison with control cells expressing GFP, ectopic expression of Fezf2 in DG lineages substantially inhibits transcription of 14% DG identity genes, including *Prox1* and *Lhx9* (Fig. 2k, l).

To investigate whether Fezf2 acts as a master transcription factor dictating CA lineage identity, cells expressing high levels of *Fezf2-shRNA* (short hairpin RNA) are enriched for transcriptome analysis. Compared with control, Fezf2 downregulation in CA lineages leads to reduced expression of 57% (48/84) CA identity genes, including *FOXG1* and *Ctip2* (Fig. 2m, n). Functionally, *Fezf2* knockout (*Fezf2^KO^*) leads to severe defects in hippocampal development, with the CA area markedly reduced at P5 (Supplementary Fig. 2j, k). Furthermore, in comparison with control, the expression levels of CA identity factors Ctip2 and FOXG1 are significantly decreased in *Fezf2^KO^* CA neurons at P5 (Supplementary Fig. 2j, l, m). Therefore, Fezf2 is indeed a hitherto poorly characterized key CA identity gene crucial for specifying CA lineage identity.

We next explored whether Fezf2 regulates the transcription of CA and DG identity genes by directly binding to these gene loci. Indeed, in both CA and DG NSCs at E13.5, Fezf2 binds to key identity genes but fails to exhibit mitotic retention in mitosis (Supplementary Fig. 3a–d). In comparison, Prox1 barely binds to lineage identity genes in CA NSCs at E13.5 (Supplementary Fig. 3a, b). Notably, our CUT&Tag analysis revealed that, in DG NSCs at E13.5, Fezf2 and Prox1 share 103 common binding peaks in 80 target genes, including key CA identity genes such as *Fezf2* itself and *Nrxn3* (Supplementary Fig. 3a, b). These results indicated that Fezf2 competes with Prox1 for binding to common peaks in lineage identity genes in DG but not CA NSCs. Strongly supporting this notion, the binding affinity of Fezf2 to lineage identity genes is significantly increased in *Prox1^Emx1-cKO^*DG cells at E14.5 (Supplementary Fig. 3e–g). Furthermore, in comparison to DG at E13.5, although Fezf2 still binds to a subset of lineage identity genes in DG at E14.5, its binding affinity is significantly reduced (Supplementary Fig. 3h–j). By contrast, the binding affinity of Prox1 to these identity genes at E14.5 is substantially increased (Supplementary Fig. 3h–j). These results demonstrate that Prox1 eventually becomes the “winner” in this Prox1 versus Fezf2 competition for identity gene binding in DG NSCs, primarily due to its ability to retain at key identity gene loci across cell divisions (Supplementary Fig. 3k). Together, Fezf2 competes with Prox1 for binding to key lineage identity genes in DG NSCs. Therefore, timely suppression of Fezf2 in DG lineages is crucial for the specification and maintenance of DG lineage identity.

Collectively, our results indicated that, Fezf2 not only promotes CA identity gene transcription and inhibits DG identity gene expression but also competes with Prox1 for identity gene binding in DG NSCs. Therefore, Prox1 and Fezf2 establish DG and CA lineage identity respectively, mainly through their mutual suppression (Fig. 2o). Fezf2 poses multiple threats to Prox1 in safeguarding DG lineage identity, explaining why Prox1 is preferentially retained at the *Fezf2* gene locus in dividing DG NSCs.

### Redundant regions in Prox1 promote its condensate formation in DG NSCs at interphase

To rigorously investigate the functional significance of Prox1 mitotic retention, we sought to introduce mutations to Prox1 that specifically compromise its mitotic retention ability in mitosis without influencing its transcriptional function in interphase. Biomolecular condensates have been implicated in a wide range of cellular processes^50–57^, including mitotic retention^16,58^. Since previous work indicated that the self-association and condensate formation ability is essential for mitotic retention of transcription factors^16,58^, we first examined whether Prox1 protein can self-associate. Indeed, FLAG-tagged Prox1 and HA-tagged Prox1 are ready to interact in coimmunoprecipitation (coIP) assays using HEK293T cell extracts (Supplementary Fig. 4a, b). Domain mapping analysis indicated that the N2 domain is both necessary and sufficient for mediating Prox1 self-association (Supplementary Fig. 4a, b). Importantly, simultaneous deletion of M1 and M2, two small motifs of 27 and 26 amino acids respectively within the N2 region, abolishes the self-association ability of Prox1 (Supplementary Fig. 4a–e). Therefore, N2M (M1 and M2), two redundant motifs in the N-terminus of Prox1, mediates its self-association.

We next investigated whether phase separation drives Prox1 condensate formation. Firstly, we performed fluorescence recovery after photobleaching (FRAP) analysis to probe the dynamics of Prox1 condensates in HEK293T cells and found that the GFP-Prox1 fluorescent signal recovers a few minutes after photobleaching (Fig. 3a, b), indicating that Prox1 condensates are dynamic. Secondly, we performed *in vivo* condensate formation assay in HEK293T cells by transiently expressing GFP-tagged N1, N2 or N3 region of Prox1. While Prox1.N1 and Prox1.N2 are ready to form discrete condensates, Prox1.N3 fails to do so (Fig. 3c–e). Consistently, Prox1.ΔN1N2 fails to form condensates in HEK293T cells (Fig. 3c–e). Thirdly, the Predictor of Natural Disordered Regions (PONDR) software predicts that Prox1.N1 and Prox1.N2 are composed of intrinsically disordered regions (IDRs) driving phase separation (Fig. 3c)^56,57^. Together, the IDRs in Prox1.N1 and N2 promote Prox1 condensate formation in interphase.

**Fig. 3.**
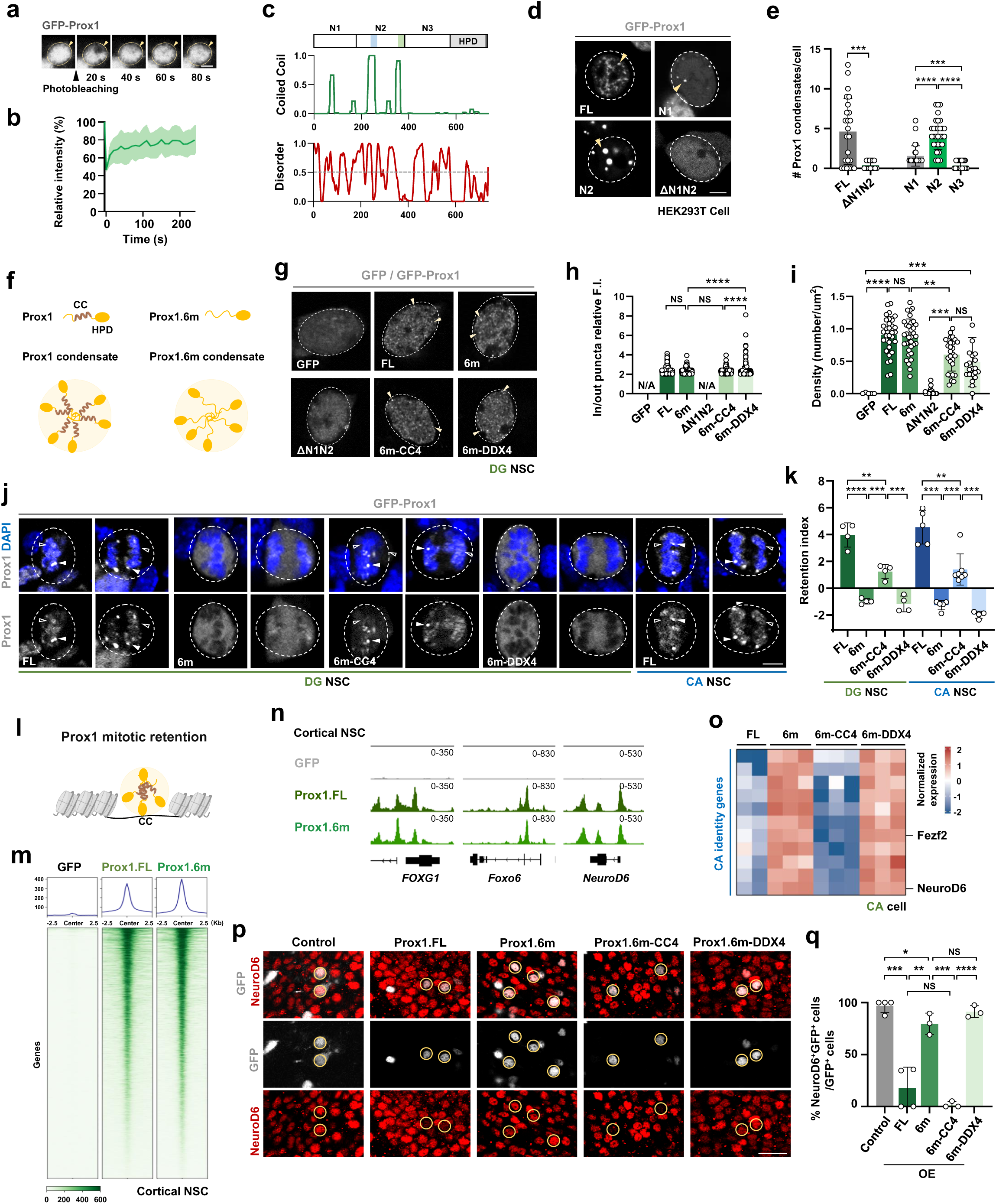
Coiled-coil-based strong condensate formation underlies Prox1 mitotic retention and CA lineage suppression in DG NSCs. (**a**, **b**) FRAP analysis of GFP-Prox1 condensates highlighted by the white boxes in HEK293T cells (**a**) and quantifications (**b**). n=8 cells. Scale bar, 1 μm. (**c**) The Prox1 protein was predicted to contain coiled-coil motifs (top) and IDRs (bottom) in the N-terminus. (**d**, **e**) Condensate formation assay of GFP-Prox1 variants in HEK293T cells (**d**) and quantifications of the condensate number per cell (**e**). n=24, 15, 27, 31, 21 cells, respectively; \*\**p* < 0.01, \*\*\**p* < 0.001,\*\*\*\**p* < 0.0001; Student’s t-test. Scale bar, 5 μm. (**f**) Schematic diagrams illustrating Prox1 protein containing CC (coiled-coil) motifs and Prox1.6m protein with CC impaired (top). IDRs in both Prox1 and Prox.6m mediate condensate formation (bottom). (**g–i**) *In vivo* condensate formation assay of GFP and GFP-Prox1 variants in DG NSCs at E14 (**g**) and quantifications of condensate fluorescent intensity (F.I.) (**h**) and condensate density (**i**). n=30, 34, 36, 31, 21 cells, respectively; N/A: not applicable; ns, not significant, \*\*\*\**p* < 0.0001; one-way ANOVA (**h**), Student’s t-test (**i**). Scale bar, 5 μm. (**j**, **k**) Representative images of metaphase and anaphase DG NSCs at E14 expressing indicated GFP-Prox1 variants, with solid and empty arrowheads indicate Prox1 foci associated with chromosomes and uniform Prox1 decorating chromosomes respectively (**j**). Quantifications of retention index for GFP-Prox1 in DG or CA NSCs (**k**). The retention index of positive values indicates enrichment on the chromosomes, whereas the retention index of negative values indicates exclusion from the chromosomes. n=4-7 cells; \*\**p* < 0.01, \*\*\*\**p* < 0.0001; Student’s t-test. Scale bar, 5 μm. (**l**) Schematic diagram illustrating Prox1 mitotic retention mediated by the CC motif. (**m**) Heat map of GFP (control), GFP-Prox1 and GFP-Prox1.6m CUT&Tag peaks in cortical NSCs. (**n**) Genomic browser image of GFP (control), GFP-Prox1 and GFP-Prox1.6m CUT&Tag tracks in cortical NSCs. (**o**) Heat map showing the expression of CA identity genes in CA lineage cells overexpressing GFP-Prox1 variants as indicated. (**p**, **q**) Representative confocal images of CA neurons overexpressing Prox1 variants stained with NeuroD6 (**p**) and quantifications of the percentage of NeuroD6^+^ cells (**q**). n = 4, 4, 3, 3, 3 animals, respectively; ns, not significant, \**p* < 0.05, \*\**p* < 0.01, \*\*\**p* < 0.001, \*\*\*\**p* < 0.0001; Student’s t-test. Scale bar, 20 μm.

### Coiled-coil-based strong condensate formation underlies Prox1 mitotic retention in DG NSCs

We next investigated whether the self-association or condensate formation ability underlies Prox1 mitotic retention in DG NSCs. AlphaFold3 predicts that the M1 and M2 motifs mediate Prox1 oligomerization (Supplementary Fig. 4f)^59^. Transiently expressed GFP-Prox1 in DG NSCs at E14 forms discrete condensates and is retained on mitotic chromosomes, resembling endogenous Prox1 (Supplementary Fig. 4g, h). In sharp contrast, self-association-defective GFP-Prox1.ΔN2M fails to retain on chromosomes of dividing DG NSCs but instead exhibits uniform cellular distribution excluding chromosomes (Supplementary Fig. 4g, h). Protein condensation is driven by either strong and specific interactions between folded domains or weak and less specific interactions mediated by IDRs^56,60,61^. Interestingly, sequence analysis of Prox1.N2M reveals that it is mainly composed of coiled-coil (CC) motifs (Fig. 3c). We therefore investigated whether restoring condensate formation ability of certain strength can rescue the mitotic retention defects of Prox1.ΔN2M. When Prox1.ΔN2M is fused with Pros.N7, the coiled-coil motif in transcription factor Prospero (Supplementary Fig. 4i)^58^, mitotic retention ability of Prox1.ΔN2M is restored (Supplementary Fig. 4g, h). By contrast, fusion with Pros.N7.5m^58^, the mutant form of N7 with its coiled-coil motif specifically impaired (Supplementary Fig. 4i), or with IDRs, such as the N-terminal region of DDX4 or FUS (Supplementary Fig. 4i)^62^, fails to restore mitotic retention of Prox1.ΔN2M in DG NSCs (Supplementary Fig. 4g, h). Therefore, strong condensate formation mediated by structured domains such as CC but not weak condensation mediated by IDRs underlies Prox1 mitotic retention in DG NSCs.

We then postulated that mutations partially impairing the condensate formation ability of Prox1 might selectively perturb its chromosome retention in mitosis without affecting its transcriptional function in interphase. We therefore mutagenized six aromatic or hydrophobic residues in the N2M region to serine residues to partially perturb the CC-based strong interaction (Supplementary Fig. 4j, k) and designated this mutated version of Prox1 as Prox1.6m. Intriguingly, whereas the condensate formation ability of Prox1.6m is comparable to Prox1 in DG NSCs at interphase (Fig. 3f–i, Supplementary Fig 4l, m), it loses its strong self-association ability and fails to retain on mitotic chromosomes of DG NSCs (Fig. 3j, k, Supplementary Fig. 4n). To more vigorously validate this notion, we fused Prox1.6m with either strongly structured CC motif such as CC4 in Kibra (Supplementary Fig. 4i)^60^, or with IDRs such as DDX4_N_. Notably, although Prox1.6m-DDX4_N_ and Prox1.6m-CC4 show comparable condensate formation ability in interphase, Prox1.6m-CC4 but not Prox1.6m.DDX4_N_ exhibits mitotic retention in DG NSCs, indicating that coiled-coil-based strong condensate formation underlies Prox1 mitotic retention in DG NSCs (Fig. 3j–l). Therefore, the 6m mutations specifically perturb the mitotic retention but not the interphase condensate formation ability of Prox1.

We next attempted to test whether the 6m mutations attenuate the chromatin-binding affinity of Prox1. To avoid the potential influence of endogenous Prox1, we performed CUT&Tag-seq in cortical NSCs, where Prox1 is barely detectable. Upon being transiently expressed in cortical NSCs for 20 h (shorter than one cell cycle) and followed by anti-GFP CUT&Tag, GFP-tagged Prox1.6m and Prox1.FL exhibit similar binding affinity genome-wide or at specific CA identity gene loci (Fig. 3m, n, Supplementary Fig. 4o). Therefore, Prox1.6m, with its interphase condensate formation ability being comparable with Prox1.FL, possesses normal chromatin-binding capacity in NSCs.

### Mitotic retention ability is crucial for Prox1 to suppress CA lineage identity

We then sought to examine whether the mitotic retention ability is critical for Prox1 to inhibit the transcription of CA identity genes. We first induced the expression of GFP-Prox1 variants in CA NSCs at E13 and assessed their mitotic retention ability. Interestingly, exogenously expressed Prox1.FL and Prox1.6m-CC4 but not Prox1.6m or Prox1.6m-DDX4_N_ exhibit mitotic retention in CA NSCs (Fig. 3j, k). We next performed transcriptome analysis to investigate whether the mitotic retention ability of Prox1 underlies the CA-to-DG neuronal identity transition induced by Prox1 overexpression. Indeed, ectopic expression of Prox1.FL and Prox1.6m-CC4 but not Prox1.6m or Prox1.6m-DDX4_N_ leads to significantly reduced transcription in key CA identity genes such as *Fezf2* and *NeuroD6* (Fig. 3o). Therefore, the mitotic retention ability is essential for Prox1 to suppress CA lineage identity.

To further validate this notion, we induced the expression of Prox1 variants in CA RGCs at E13 and examined the expression levels of CA identity factor NeuroD6 at E15. While mitotic retention-competent Prox1.FL and Prox1.6m-CC4 effectively suppress the expression of NeuroD6, mitotic retention-deficient Prox1.6m or Prox1.6m-DDX4_N_ exhibit minimal effects on NeuroD6 expression (Fig. 3p, q). Together, our results revealed that Prox1 mitotic retention is both necessary and sufficient for suppressing the alternative CA lineage identity.

### Prox1 suppresses CA lineage identity via PRC2 recruitment and H3K27me3 deposition

We next sought to understand by what mechanism Prox1 inhibits the transcription of CA identity genes and why mitotic retention ability is essential for Prox1 to execute this critical task. First, we performed *in vivo* CUT&Tag with DG NSCs at E13.5 for H3K9me3 and H3K27me3, two major repressive histone marks. Notably, H3K27me3 and Prox1 display remarkable co-occupancy at a large subset of CA identity gene loci in DG cells (Fig. 4a, b), suggesting that Prox1 inhibits CA identity gene transcription via promoting local H3K27me3 deposition. Significantly, these Prox1-bound CA loci with H3K27me3 marks show enrichment of EZH2 and SUZ12, core subunits of PRC2 (Fig. 4c, d), strongly hinting that Prox1 recruits PRC2 to catalyze H3K27me3 deposition in DG NSCs. In accordance with this notion, CA identity gene loci in CA NSCs show limited enrichment of Prox1 binding or H3K27me3 marks (Fig. 4e). More importantly, H3K27me3 deposition at these CA loci are substantially reduced in *Prox1^Nes-cKO^*DG neurons (Fig. 4f, Supplementary Fig. 5a). Furthermore, ectopic expression of Prox1 in CA lineages led to markedly increased H3K27me3 deposition at CA identity gene loci (Fig. 4f, Supplementary Fig. 5b). Taken together, Prox1 is both necessary and sufficient to induce H3K27me3 deposition at CA identity gene loci, presumably via PRC2 recruitment.

**Fig. 4.**
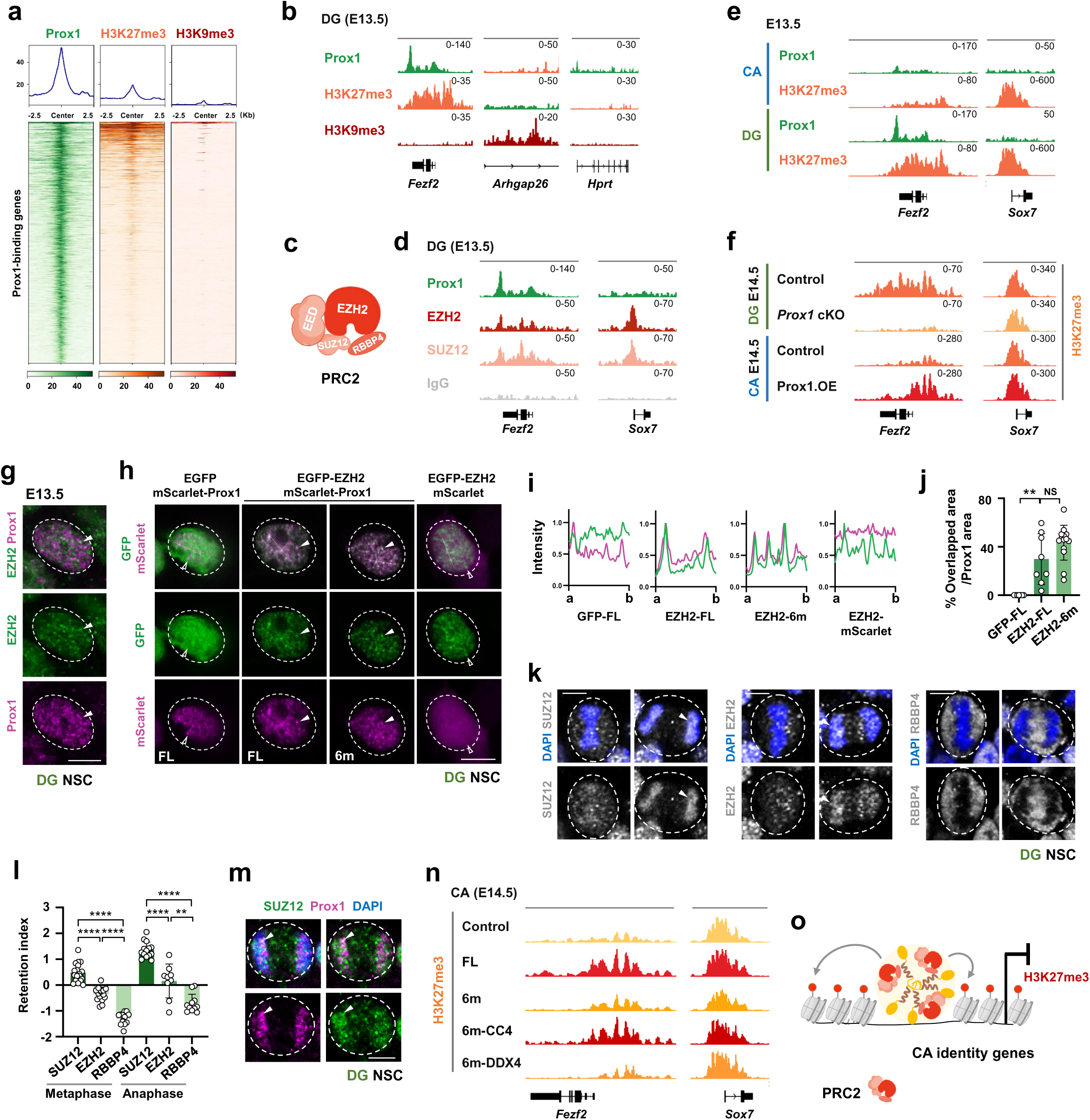
Prox1 suppresses CA lineage identity via PRC2 recruitment and H3K27me3 deposition. (**a**) Heat maps of Prox1, H3K27me3 and H3K9me3 CUT&Tag peaks in DG NSCs at E13.5. (**b**) Genome browser images of CUT&Tag for Prox1, H3K27me3 and H3K9me3 tracks at *Fezf2*, *Arhgap26* and *Hprt* gene locus in DG NSCs at E13.5. (**c**) Schematic diagram showing the core subunits of PRC2 complex. (**d**) Genome browser images of CUT&Tag for Prox1, EZH2, SUZ12 and IgG tracks at *Fezf2* and *Sox7* gene locus in DG NSCs at E13.5. (**e**) Genome browser images of CUT&Tag for Prox1 and H3K27me3 tracks at *Fezf2* and *Sox7* gene locus in CA (top) and DG (bottom) NSCs at E13.5. (**f**) Genome browser images of CUT&Tag for H3K27me3 tracks at *Fezf2* and *Sox7* gene locus in control and *Prox1^Nes-cKO^* DG cells at E14.5 (top) and in control and Prox1-overexpression CA cells at E14.5 (bottom) respectively. (**g–j**) Representative images of WT DG NSCs at E13.5 stained for Prox1 and EZH2 (**g**) and DG NSCs at E14 co-expressing GFP, GFP-EZH2, mSarlet or mSarlet-Prox1 variants (**h**), line plots of fluorescent intensity (I) and quantifications of overlapped area (**j**). n=7, 8, 14 cells, respectively; **p < 0.01, NS, not significant; Student’s t-test. Scale bar, 5 μm. (**k**, **l**) Representative images of metaphase and anaphase DG NSCs at E13.5 stained for SUZ12 (left), EZH2 (middle), RBBP4 (right) and DAPI (**k**) and quantifications of retention index (**l**). The retention index of positive values indicates enrichment on the chromosomes, whereas the retention index of negative values indicates exclusion from the chromosomes. n=18, 14, 15, 11, 13, 11 cells respectively; **p < 0.01, ****p < 0.0001; Student’s t-test. Scale bar, 5 μm. (**m**) Representative imaging showing E13.5 DG NSCs at anaphase stained for SUZ12, Prox1 and DAPI. Scale bar, 5 μm. (**n**) Genome browser images of CUT&Tag for H3K27me3 tracks at *Fezf2* and *Sox7* gene locus in CA cells at E14.5 overexpressing GFP (control) or GFP-Prox1 variants as indicated. (**o**) Schematic diagram showing Prox1 and PRC2 form co-condensates and inhibit transcription of CA identity genes via H3K27me3 deposition.

To investigate whether Prox1 and PRC2 form a protein complex, we carried out coIP experiments in HEK293T cells. FLAG-tagged N1, N2 but not N3 region of Prox1 interact with EGFP-EZH2 (Supplementary Fig. 5c, d). Furthermore, FLAG-Prox1 exhibits specific interaction with EGFP-SUZ12 (Supplementary Fig. 5e). Therefore, Prox1 recruits PRC2 through its specific interaction with PRC2 core subunits to induce H3K27me3 deposition at CA identity gene loci (Supplementary Fig. 5f). In accordance, previous studies demonstrated that conditional knockout of PRC2 subunit EED leads to specific and drastic defects in DG development^63^.

We next asked whether Prox1 colocalizes with PRC2 in DG NSCs. Both Prox1 and EZH2 form small-sized discrete condensates and partially colocalize in E13.5 DG NSCs at interphase (Fig. 4g). We then investigated whether Prox1 can recruit PRC2 into its condensates in E14 DG NSCs at interphase. Notably, mScarlet-Prox1 forms discrete nuclear condensates by itself and co-condensates with GFP-EZH2 but not EGFP upon being co-expressed in DG NSCs (Fig. 4h–j). Consistently, although GFP-EZH2 forms nuclear condensates in DG NSCs, it fails to partition mScarlet into its condensates (Fig. 4h–j). Importantly, Prox1.6m, similar to Prox1.FL, effectively recruits EZH2 into its condensates (Fig. 4h–j, Supplementary Fig. 5f, g), further supporting the notion that the transcriptional function of Prox1.6m is comparable to Prox1.FL in DG NSCs at interphase.

Given that Prox1 exhibits mitotic retention in DG NSCs, we next explored whether PRC2 core subunits also remain associated with mitotic chromosomes. Remarkably, SUZ12, EZH2 but not RBBP4 exhibit retention on mitotic chromosomes of DG NSCs (Fig. 4k, l), strongly suggesting that a Prox1-PRC2 complex containing a subset of PRC2 subunits is mitotically retained in DG NSCs. Indeed, Prox1 and SUZ12 form co-condensates and retain on chromosomes of E13.5 DG NSCs at anaphase (Fig. 4m).

Finally, to assess whether the mitotic retention ability is important for Prox1 to promote H3K27me3 deposition on key CA identity gene loci, we expressed Prox1 variants in CA lineages 36 h (1.5 cell cycles) and carried out anti-H3K27me3 CUT&Tag analysis. For *Fezf2* locus, substantially higher levels of H3K27me3 deposition are induced by Prox1.FL or Prox1.6m-CC4 than it is by Prox1.6m or Prox1.6m-DDX4_N_ in CA lineages (Fig. 4n), highlighting the importance of Prox1 mitotic retention for epigenetic silencing of key CA identity genes in a timely and precise manner (Fig. 4o).

### Mitotic bookmarking by Prox1 is crucial for hippocampal development in mice

Considering that Prox1.6m exhibits normal condensate formation, chromatin-binding and PRC2 recruitment ability in interphase NSCs yet specifically impaired mitotic retention ability in dividing DG NSCs, we generated *Prox1.6m* knock-in (referred to as *Prox1.6m^KI^*) mice to investigate the functional significance of Prox1 mitotic retention under physiological conditions (Fig. 5a). Strikingly, all heterozygous *Prox1.6m^KI^* mice that we obtained died a few days after birth (Fig. 5b), demonstrating that mitotic retention of Prox1 is critical for mouse early development and survival.

**Fig. 5.**
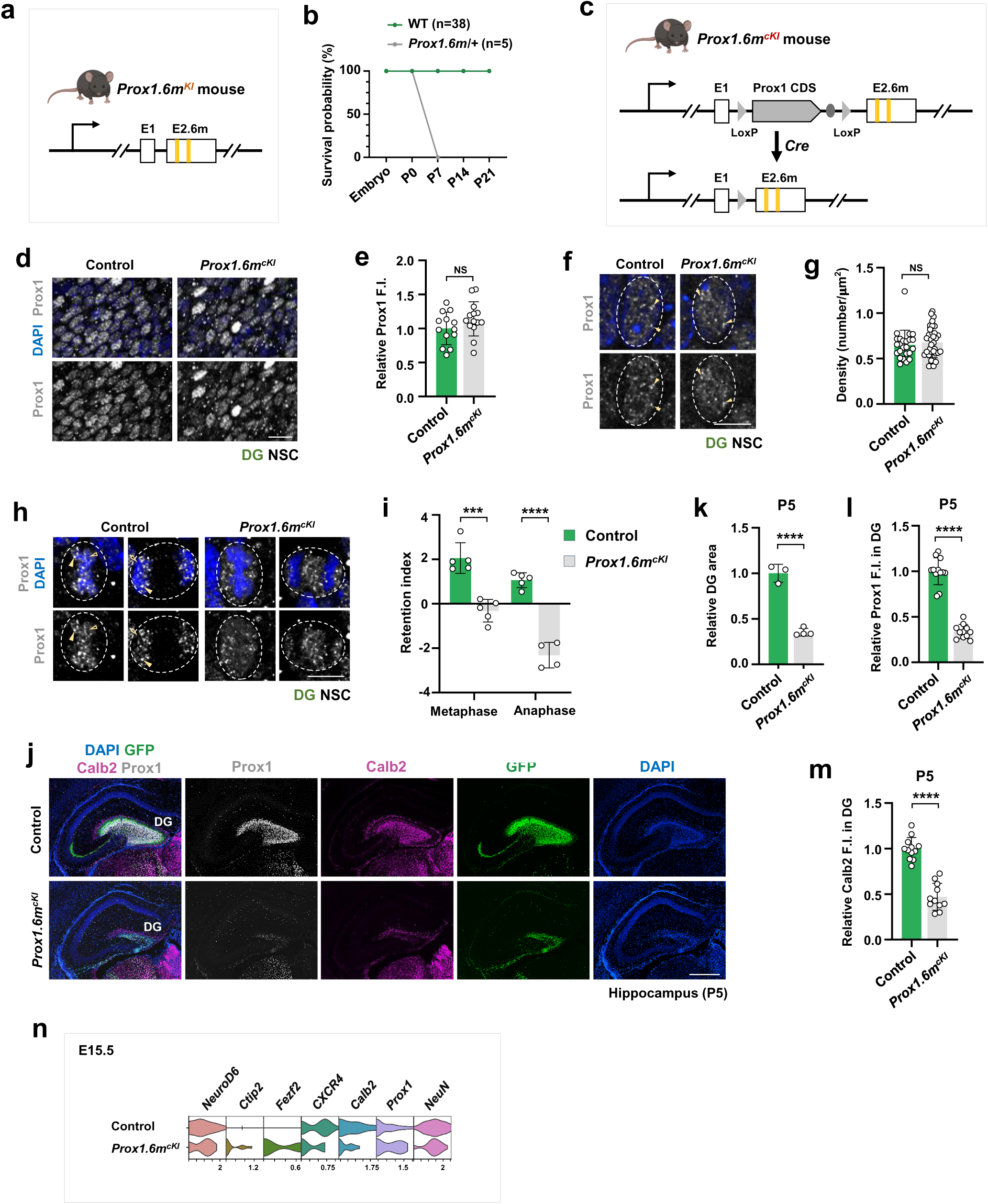
Mitotic bookmarking by Prox1 is crucial for hippocampal development in mice. (**a**) The strategy for generating *Prox1.6m* knockin (KI) mice. (**b**) The survival curve of WT and *Prox1.6m^KI^* mice. (**c**) The flowchart of creating *Prox1.6m* conditional knockin (cKI) mice. (**d**, **e**) Representative images of control (*Emx1-Cre; Prox1.^fl/+^*) and Prox1.6mcKI (*Emx1-Cre; Prox1^fl/6m.fl^*) DG at E13 stained for Prox1 (**d**) and quantifications of relative Prox1 expression levels (**e**). n=13, 14 brain sections, respectively; NS, not significant; Student’s t-test. Scale bar, 20 μm. (**f**, **g**) Sample images of control and *Prox1.6m^cKI^* DG NSCs at E13 stained for Prox1 (**f**) and quantifications of condensate density (**g**). n=24, 35 cells, respectively; NS, not significant; Student’s t-test. Scale bar, 5 μm. (**h**, **i**) Representative images of control and *Prox1.6m^cKI^* metaphase and anaphase DG NSCs at E13 stained for Prox1 (**h**) and quantifications of retention index (**i**). n=5, 5, 5, 4 cells, respectively; ***p < 0.001, ****p < 0.0001; Student’s t-test. Scale bar, 5 μm. (**j–m**) Representative images of control and *Prox1.6m^cKI^* hippocampus at P5 stained for Prox1, Calb2 and DAPI (**j**) and quantifications of relative DG area (**k**; n=3, 4 animals), relative Prox1 (**l**; n=12 brain sections) and Calb2 (**m**; n=11 brain sections) fluorescent intensity. ****p < 0.0001; Student’s t-test. Scale bar, 500 μm. (**n**) Violin plots showing expression of lineage identity genes in control and *Prox1.6m^cKI^* DG neurons.

We therefore generated NSC lineage-specific *Prox1.6m* knock-in mice (*Emx1*-Cre; *Prox1^fl^*^/*6m.fl*^; referred to as *Prox1.6m^.cKI^*) (Fig. 5c, Supplementary Fig. 6a). In comparison to littermate control (*Emx1*-Cre; *Prox1^fl^*^/+^), *Prox1.6m^.cKI^* exhibits comparable Prox1 expression levels and similar condensate formation capacity in interphase but fails to retain on chromosomes of DG NSCs (Fig. 5d–i).

Remarkably, the DG areas in *Prox1.6m^.cKI^* mice at P5 are significantly smaller than they are in the littermate control, with the expression levels of DG identity marker Calb2 drastically reduced (Fig. 5j–m), highlighting the functional significance of Prox1 mitotic bookmarking for safeguarding DG lineage identity and DG development.

To investigate whether defective Prox1 mitotic retention in *Prox1.6m^.cKI^*leads to ectopic expression of *Fezf2* in DG lineages, we performed scRNA-seq using control and *Prox1.6m^.cKI^* hippocampus at E15.5. Indeed, violin plots revealed that key CA identity genes such as *Fezf2* and *Ctip2* are barely detectable in control DG neurons but are ectopically expressed in *Prox1.6m^.cKI^* DG neurons (Fig. 5n, Supplementary Fig. 6b). Therefore, mitotic bookmarking by Prox1 is crucial for preserving DG lineage identity memory.

### Histone bivalency at *Fezf2* locus necessitates Prox1 mitotic bookmarking in DG NSCs

We next asked why the timely silencing of *Fezf2* in DG lineages requires mitotic retention of Prox1. While examining the deposition of various histone marks, we unexpectedly noted that a subset of CA identity genes, including *Fezf2*, *FOXG1* and *NeuroD2*, are bivalent genes, with each locus carrying both the active H3K4me3 mark and the repressive H3K27me3 mark in DG NSCs (Fig. 6a, b). This means that reduction of the H3K27me3 levels below a certain threshold could lead to immediate derepression of CA identity genes and lineage identity crisis. Intriguingly, in comparison to other bivalent CA identity genes, the H3K4me3 to H3K27me3 deposition ratio is much higher at *Fezf2* locus (Fig. 6b), rendering it one of the most dangerous CA identity gene loci. This further explains why *Fezf2* is the only key CA identity gene locus that is mitotically bookmarked by Prox1.

**Fig. 6.**
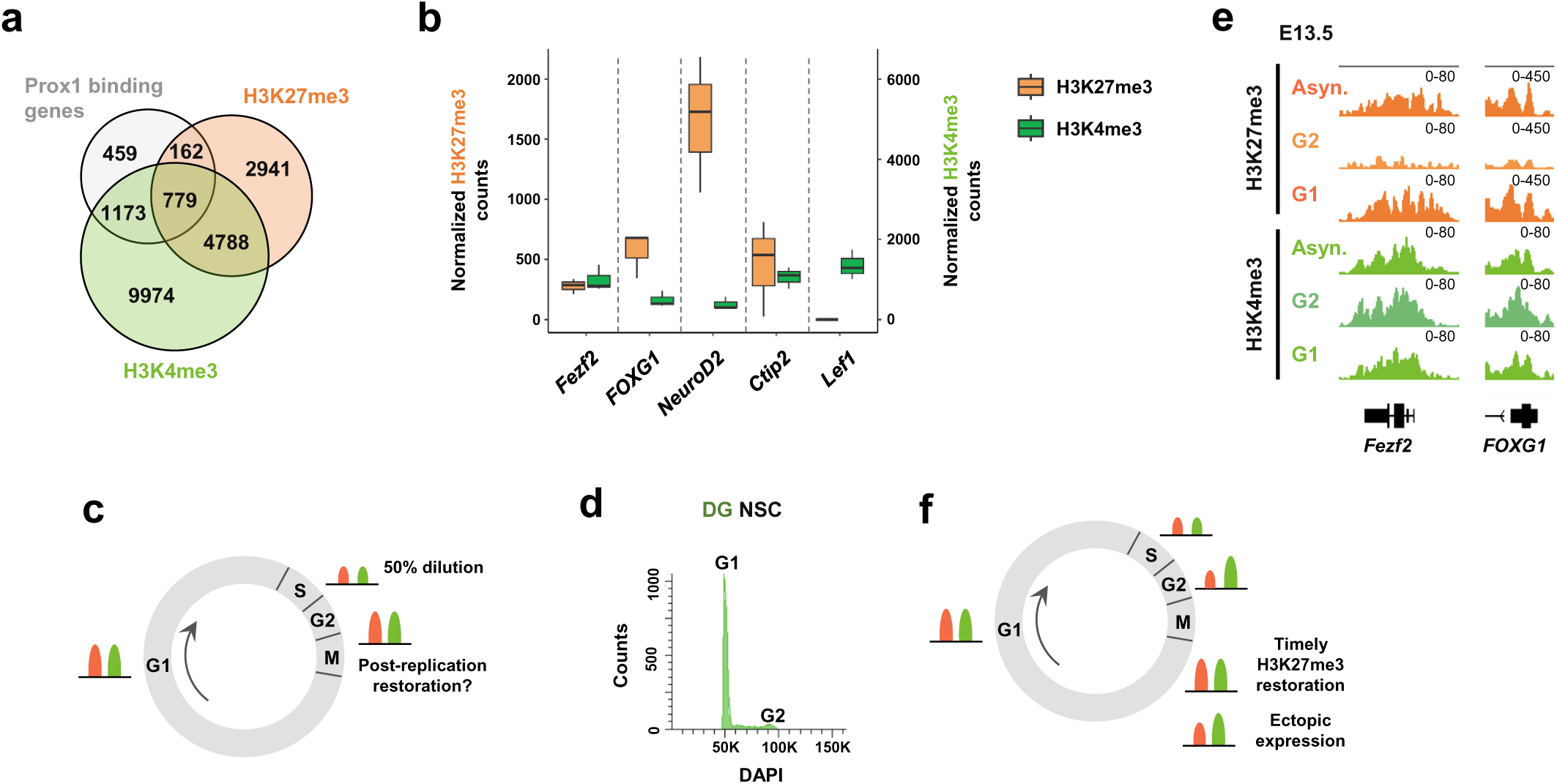
Histone bivalency at *Fezf2* locus necessitates Prox1 mitotic bookmarking in DG NSCs. (**a**) Venn diagram showing the overlap of Prox1 binding genes and genes marked by H3K27me3 and H3K4me3. (**b**) Box plots demonstrating that *Fezf2* gene has a higher H3K4me3:H3K27me3 ratio than other bivalent CA identity genes. The non-bivalent DG identity gene *Lef1* serves here as a control. (**c**) Schematic diagram showing the levels of histone marks are diluted to half after DNA replication at S phase and restored at G2 and G1 phase. (**d**) Purification of G1 and G2 DG NSCs after DNA content-based FACS sorting. (**e**) Genome browser examples of CUT&Tag for H3K27me3 (top) and H3K4me3 (bottom) tracks at *Fezf2* and *FOXG1* gene locus in asynchronous, G2 and G1 DG NSCs at E13.5. (**f**) Schematic diagram showing that timely H3K27me3 restoration at early G1 prevents ectopic identity gene expression, whereas slow H3K27me3 restoration at G1 results in ectopic identity gene expression.

Both enzymatic erasure and replicational dilution contribute to a reduction in H3K27me3 levels at a specific genomic locus^64^. Since H3K27me3 demethylase KDM6B is barely detectable in embryonic hippocampus^65^, the reduction in H3K27me3 levels is primarily due to replicational dilution^64,66,67^. That is, during DNA replication, due to the incorporation of newly-assembled unmodified nucleosomes, the levels of all covalent histone marks are diluted by 50% (Fig. 6c). Previous studies indicated that, after replicational dilution, different histone marks are restored at distinct speed^64,66,68^. We therefore investigated H3K4me3 vs. H3K27me3 levels in DG NSCs at different cell cycle stages. E13.5 DG NSCs at G1 or G2 stages or asynchronous DG NSCs were purified for CUT&Tag analysis (Fig. 6d). Interestingly, in comparison with asynchronous DG NSCs, H3K27me3 levels enriched at *Fezf2* locus is substantially lower at G2 and becomes comparable at G1 (Fig. 6e, f, Supplementary Fig. 6c), indicating that H3K27me3 restoration post DNA replication is relatively slow and cannot be fully restored until G1. In sharp contrast, H3K4me3 levels at *Fezf2* locus is fully restored at G2 (Fig. 6e, f, Supplementary Fig. 6d). Considering that the restoration of H3K27me3 requires its writer PRC2 and that H3K4me3 inhibits PRC2 enzymatic activity^69^, it is reasonable that mitotic bookmarking by Prox1 and hence timely recruitment of PRC2 to *Fezf2* locus in early G1 are crucial for preventing ectopic expression of *Fezf2* and lineage identity crisis.

## Discussion

The human brain is composed of neuronal lineages of enormous diversity. However, how these diverse neuronal lineage identities are established in brain development remains not fully understood. Even less is known about how specific neuronal lineage identity memory is faithfully transmitted from neural stem cells to their progenies across cell generations. Our results identified the transcription factor Prox1 as a *bona fide* mitotic bookmark in mouse hippocampal development and revealed mitotic bookmarking as a fundamental strategy for preserving neuronal lineage identity memory in the developing mammalian brains.

One major obstacle to the identification of *bona fide* mitotic bookmarks mammalian brain development is the challenge to isolate limited number of mitotic neural progenitors, followed by low-input omics approaches. Here, based on our newly-developed BISMIB (<u>bi</u>nding <u>s</u>ites of <u>mi</u>totic <u>b</u>ookmark identification) method for isolating mitotic NSCs from developing fly brains^15^, we designed a new pipeline suitable for enriching mitotic and interphase NSCs from developing mouse brains. With this new pipeline, we successfully identified key CA identity gene *Fezf2* as one critical gene locus mitotically bound by Prox1 in DG NSCs of embryonic hippocampus. This new pipeline promises to facilitate the identification of gene loci mitotically bound by other bookmarking factors in diverse mammalian tissues or organs.

A major challenge in understanding the functional significance of mitotic bookmarking in the developing brains is to disentangle the function of mitotic bookmarks in mitosis versus interphase. Previous neatly-designed strategies, including the mitosis-specific degradation (MD) system and the auxin-inducible degradation (AID) system^10,73^, although work very effectively in cultured cells, are not applicable for studies in the developing tissues or organs^15^. Here, we adopted a different strategy by generating a mutant version of Prox1 that is specifically defective in mitotic retention, with its functional properties in interphase, including its expression levels, transcriptional condensate formation, recruitment of the PRC2 complex and chromatin binding abilities, remaining intact. Such a retention-deficient version of Prox1 allowed us to assess the functional importance of Prox1 mitotic bookmarking *per se* in mouse hippocampal development. Strikingly, the *Prox1* retention-deficient knock-in mice *Prox1.6m^KI^*, with their mitotic retention ability specifically compromised, are heterozygous lethal a few days after birth, highlighting the physiological significance of Prox1 mitotic retention for mouse early development. *Prox1^KO/+^* mice have been found to die within 2 to 3 days after birth, presumably due to haploinsufficiency of Prox1 in the normal development of the lymphatic system^74^. Therefore, Prox1 mitotic bookmarking is very likely to plays crucial roles in multiple organs in mouse early development. More importantly, NSC lineage-specific *Prox1.6m^cKI^* mice exhibit drastic DG developmental defects, pinpointing the functional importance of Prox1 mitotic bookmarking in hippocampal early development.

Mechanistically, we found that the unique mitotic bookmarking ability is critical for Prox1 to “win” two “battles” for safeguarding DG neuronal lineage identity. The first battle is between Prox1 and Fezf2 in competitive binding to key CA identity gene loci. Within at least a subset of DG NSCs at E13.5, Prox1 and Fezf2 bind to key CA identity gene loci such as *Fezf2* with comparable affinity. The long residence time of Prox1 at *Fezf2* locus throughout cell cycle, including mitosis, precludes the competitive binding of Fezf2 to the same locus in early G1, ensuring transcriptional repression of *Fezf2* by Prox1 in DG NSCs (Supplementary Fig. 3k). In accordance, at E14.5, one cell cycle after E13.5, Prox1 and Fezf2 exhibit increased and decreased binding to *Fezf2* locus respectively (Supplementary Fig. 3i, j). The second battle is between H3K27me3 and H3K4me3 deposition on the bivalent key CA identity gene locus *Fezf2* for maintaining DG lineage identity across cell generations. Mitotic retention of Prox1-PRC2 complex at *Fezf2* locus promotes timely and precise H3K27me3 restoration after replicational dilution, therefore preventing H3K27me3 deposition being outcompeted by H3K4me3 and hence ectopic expression of CA identity genes in DG NSCs (Fig. 6e, f). Together, mitotic bookmarking by Prox1 preserves DG neuronal lineage identity memory via promoting timely H3K27me3 restoration (Fig. 7).

**Fig. 7.**
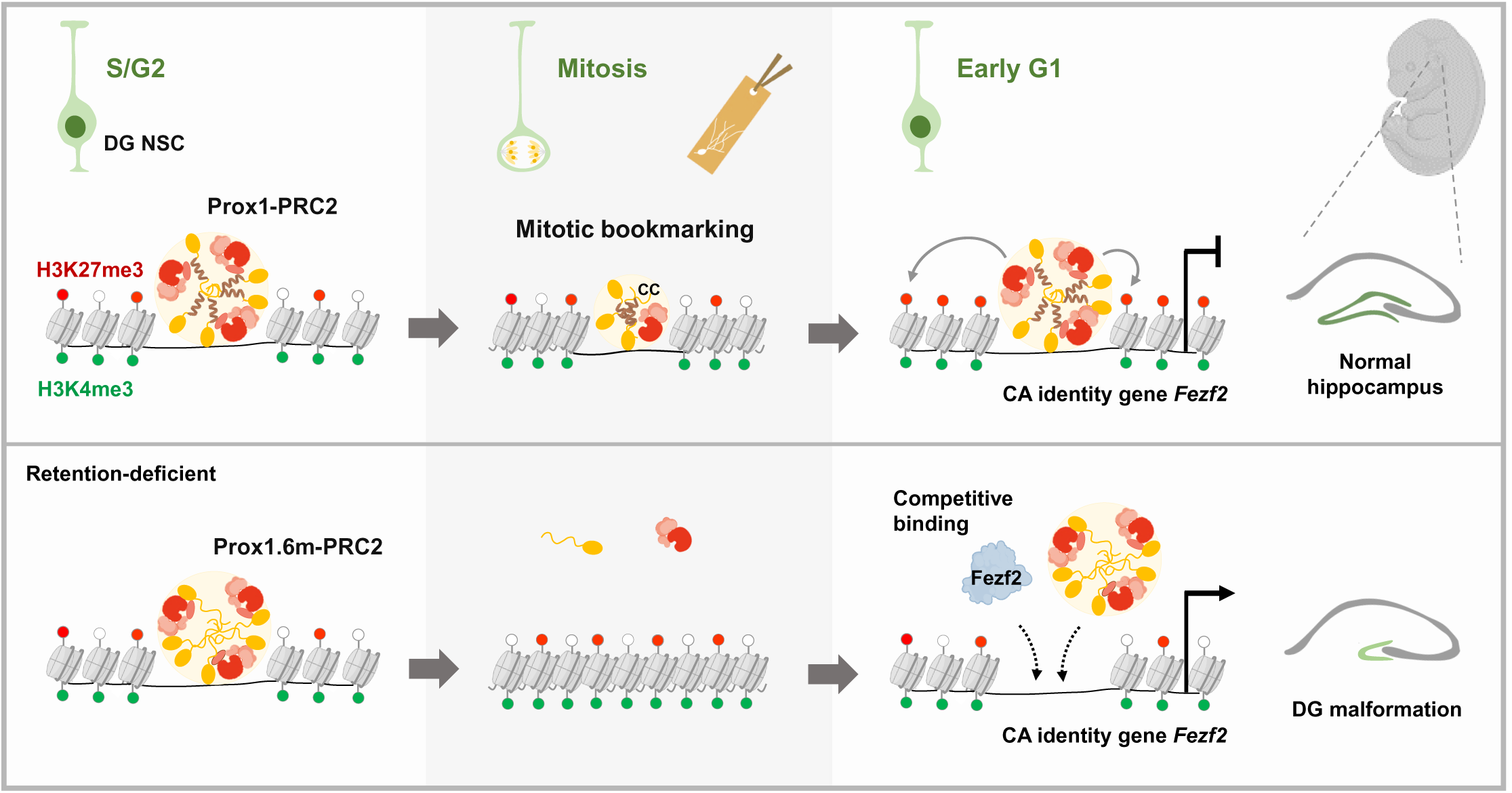
Mitotic bookmarking by Prox1 preserves hippocampal neuronal lineage identity memory via timely H3K27me3 restoration. A schematic model depicting that mitotic bookmarking by Prox1 at key bivalent CA identity gene *Fezf2* promotes timely and precise restoration of H3K27me3 deposition to avoid ectopic expression of CA identity genes and DG developmental defects. Coiled-coil (CC)-based strong condensate formation underlies Prox1 mitotic retention in DG NSCs.

Cross-repression between key TFs has been considered a paradigm for translating morphogen gradient into distinct neuronal identities^5,75,76^. Here we propose that mitotic bookmarking acts as a crucial component of this paradigm, rendering alternative neuronal identities to be specified in a more deterministic manner. More importantly, our results revealed that mitotic bookmarking represents a powerful strategy for ensuring faithful transmission of specific neuronal lineage identity memory across mitosis. This newly uncovered molecular paradigm is very likely to represent a fundamental principle underlying the preservation of epigenetic memory in diverse mammalian neuronal lineages. Identification and characterization of more mitotic bookmarking factors in distinct regions of the developing brains promise to not only shed fresh light on fundamental principles underlying brain patterning but also promote neurodevelopmental disorder therapies and regenerative medicine^1,37^.

## Methods

### Animals

All animal procedures used in this study were performed in accordance with protocols approved by the Institutional Animal Care and Use Committee (IACUC) of Peking University Laboratory Animal Center and were approved by the Peking University Animal Care and Use Committee (LSC-SongY-2). Animals were housed in a 12 h light/12 h dark cycle in a temperature controlled (22 °C), 50% humidity and air-circulating cabinet and provided with food and water as libitum. *Prox1^flox^* mouse line^41^ was a generous gift from Dr. Gord Fishell’s lab at Havard University and Dr. Fumio Matsuzaki’s lab at RIKEN BDR. *Prox.6m^KI^* mouse line was generated by mutagenizing six hydrophobic/aromatic residues into serine residues (6m) in the exon 2 of *Prox1* via CRISPR-Cas9 (Biocytogen; Fig. 5a, Supplementary Fig. 4j). *Prox1.6m^.fl^* mouse line was generated by introducing *Prox1* coding region sequence (CDS) flanked by loxP sites intro the first intron and mutagenizing six hydrophobic/aromatic residues into serine residues (6m) in the exon 2 of *Prox1* locus (Biocytogen; Fig. 5c, Supplementary Fig. 4j). *Fezf2^KO^* was purchased from GemPharmatech (Strain # T002633). *Nestin-CreER^T2^* was a generous gift form Dr. Songhai Shi’s lab at Tsinghua University. *Emx1-Cre* (JAX 005628) was a generous gift from Dr. Chenglin Miao’s lab at Peking University. The toe clipping of mice were incubated in Lysis Buffer containing Proteinase K (Vazyme, PD101) at 55℃ for 20 min and subsequently at 95℃ for 5 min to isolate the genomic DNA. The timed pregnancy was determined by identifying a vaginal plug (E0.5).

### Cell lines

HEK293T cells were cultured in DMEM supplemented with 10% (v/v) fetal bovine serum and 1% non-essential amino acids at 37°C in a 5% CO_2_ incubator.

### Molecular biology

For *In utero* electroporation (IUE) experiments, mScarlet-Prox1, mScarlet-Prox1.6m, EGFP-Fezf2, EGFP-Pbx1, EGFP-EZH2 or full-length, truncated or mutated versions of EGFP-Prox1 were cloned into the pCAG vector. For restoration of condensation ability, DNA fragments encoding human hnRNPA1_C_ (aa 186-320), DDX4_N_ (aa 1-236), FUS_N_ (aa 1-214), CC4 (coiled-coil motif of human Kibra; aa 359-432), fly Pros.N7 or Pros.N7.5m were added to the C-terminus of GFP-Prox1.6m or GFP-Prox1.ΔN2M via Gibson assembly. For *Fezf2* knockdown, scramble *shRNA* or *shRNA* against Fezf2 were cloned into pSicoR vector. For coimmunoprecipitation experiments, EZH2, SUZ12 and full-length or truncated versions of Prox1 were cloned into the pcDNA3.1 vector respectively. For OptoDroplet assay, the M1 (aa 230-256) and M2 (aa 346-371) fragments in Prox1.N2 were ligated together before being clone in pHR-mCherry-Cry2 plasmid.

### Tamoxifen injection

A stock solution of tamoxifen at a concentration of 20 mg/mL was prepared by dissolving tamoxifen powder (Sigma, T5648) in corn oil (Sigma, C8267) at 37°C with continuous vortexing until complete dissolution was achieved. On embryonic day 13.5 (E13.5), the tamoxifen solution was administered via intraperitoneal injection to timed-pregnant mice. The injected mice were analyzed at embryonic day 18.5 (E18.5), postnatal day 0 (P0) or postnatal day 120 (P120) respectively.

### *In utero* electroporation

Electroporation at hippocampal region was performed as previously described^83^. Timed natural matings were employed, designating noon on the day of plug identification as embryonic day 0.5 (E0.5). In brief, timed pregnant mice (E13) were deeply anaesthetized with isoflurane (Sigma, T48402), and the uterine horns were exposed through a midline incision. IUE was performed with 50 ms, 33 V unipolar pulses (BTX, ECM830) using 3mm electrodes (BTX). Plasmids were injected at concentrations ranging from 1.5 to 2.5 µg/µl, with a total volume of 0.5 µl per embryo. Embryos were subsequently harvested at different time points: 18 h post-electroporation for mitotic retention assays, 36 h post-electroporation (E14.5) for RNA sequencing and anti-H3K27me3 CUT&Tag analysis, and 48 h post-electroporation (E15) for functional assays.

### Morris water maze

Spatial learning was assessed using a water maze task. The apparatus consisted of a circular pool (120 cm diameter) filled with opaque water, featuring prominent visual cues around its perimeter. A fixed, black escape platform (10 cm diameter) was used. The procedure was conducted over two sequential phases: during the visible platform phase (days 1-2), the platform was raised 2 cm above the water line; during the subsequent hidden platform phase (days 3-6), it was submerged 2 cm below the surface. Each day, mice underwent four trials from unique starting points. They were given 60 s to locate the platform. If unsuccessful, they were gently guided to it. Upon finding the platform, mice remained there for 10 s to facilitate spatial learning. Between trials, mice were dried and placed in a holding cage for a 20-min inter-trial interval. Performance was measured by the latency to find the platform and swimming trajectories, which were automatically tracked and analyzed using the SMART 3.0 system.

### Immunohistochemistry

Animals were transcardially perfused with ice-cold 1x PBS, followed by perfusion with ice-cold 4% paraformaldehyde (PFA) for mice older than E15.5. Brains were then fixed overnight in 4% PFA at 4°C. After fixation, brains were washed three times in 1x PBS and subsequently cryoprotected by sequential immersion in 20% and 30% sucrose solutions until the tissues sank to the bottom. Brains were then embedded in O.C.T. compound (Sakura) and sectioned at 10-40 µm thickness using a cryostat (Leica, CM3050S).

After sectioning, brain slices were washed three times in PBS to remove the O.C.T. compound. Sections were then permeabilized with a solution of PBS containing 0.3% Triton X-100 for 30 minutes, followed by blocking with a solution of PBS containing 0.3% Triton X-100 and 5% donkey serum. Sections were incubated with primary antibodies diluted in blocking solution overnight at 4°C on a shaker. Antibodies used in this study include goat anti-Prox1 (1:50, R&D Systems, AF2727), goat anti-Sox2 (1:200, R&D Systems, AF2018), rabbit anti-Prox1 (1:200, Abcam, Ab101851), rat anti-Tbr2 (1:200, Thermo Fisher Scientific, 14-4875-82), rabbit anti-NeuroD1 (1:200, Abcam, Ab213725), rabbit anti-NeuN (1:200, Abcam, Ab177487), rabbit anti-NeuroD2 (1:200, Abcam, Ab104430), rabbit anti-NeuroD6 (1:200, Abcam, Ab85824), rabbit anti-FOXG1 (1:200, Abcam, Ab18259), rat anti-Ctip2 (1:200, Abcam, Ab18465), rabbit anti-Calb2 (Calretinin) (1:200, Abcam, Ab244299) and rabbit anti-Calb1 (Calbindin) (1:200, Abcam, Ab108404). Following incubation, brain sections were washed four times in PBS and incubated in blocking solution for 30 min, before being incubated with secondary antibodies and DAPI (Roche) in blocking solution for 1 h at room temperature. After washing with PBS containing 0.01% Triton X-100 for 6–8 times, the sections were mounted with antifade mounting medium. Fluorescent images were acquired using a Leica SP8 confocal microscope or a Nikon AXR laser scanning microscope.

### Dissociation of mouse embryonic NSCs, neural progenitors or neurons

Mouse brain regions were dissected in serum-free DMEM and transferred to a dissociation solution containing 2% papain in serum-free DMEM. The tissues were then gently triturated using a pipette tip to facilitate mechanical dissociation into small pieces. The resulting mixture was subsequently incubated at 37°C for 60-70 minutes. Following digestion, the reaction was terminated by adding an equal volume of DMEM complete medium containing 10% fetal bovine serum. The cell suspension was then passed through a 30 μm mesh filter to remove undigested debris.

### Condensate formation assay in DG NSCs

After immunofluorescence staining of the brain sections (10 µm thick) on coverslips, images were acquired using a Nikon AXR laser scanning confocal microscope. Image stacks were then deconvolved using the Richardson-Lucy algorithm. Puncta analysis was performed in FIJI/ImageJ, with puncta identified based on the following criteria: 1) an intensity threshold set at twice the mean intensity of the individual cell; 2) a minimum Feret’s diameter of 100 nm; and 3) a circularity index of ≥ 0.8.

The relative fluorescence intensity (F.I.) of puncta for Prox1 variants was quantified using the ratio of in-puncta intensity (mean intensity within puncta) to out-puncta intensity (mean intensity in the nucleoplasm).

Colocalization of puncta across fluorescent channels was quantified using the JACoP plugin in FIJI. Puncta areas were defined by the intensity threshold in each channel. The extent of overlap was measured using Manders’ Colocalization Coefficients (MCC).

### Condensate formation assay in HEK293T cells

HEK293T cells were plated 24 h prior to transfection in 4-chamber glass bottom dish (Cellvis). After transfection, the dishes were cultured in 37°C incubator for 20-24 h. Imaging was performed on Leica SP8 confocal. Puncta analysis was performed using criteria consistent with the previous method, employing identical thresholds for intensity and circularity (≥ 0.8), but with an adjusted minimum Feret’s diameter of 350 nm.

### RNA-seq

RNA-seq analysis was performed according to the manufacturer’s protocol (N711, Vazyme). Briefly, after digesting the mouse DG or CA neuroepithelium at indicated developmental stages, the cell pellet (50-2,000 cells, identical number of cells were used for control and experimental groups) was resuspended in a 1 µL sample buffer, with nuclease-free water being added to a final volume of 6 µL, and incubated at room temperature for 5 min. Oligo dT Primer and dNTP Mix were then added to the sample and incubated at 72℃ for 3 min. The sample was incubated on ice for 2 min before FS Buffer V2, DTT, RNase Inhibitor, Discover-sc TS Oligo V2, Discover-sc Reverse Transcriptase and Nuclease-free distilled H_2_O were added to a total of 20 µL. The mixture was amplified for the first strand cDNA, before the Discover-sc WTA PCR Primer, Discover-sc PCR Mix and Nuclease-free H_2_O were added to the sample for full-length cDNA amplification for 15 cycles. Product purification was performed by using VAHTS DNA Clean Beads (N411, Vazyme). After washes, the beads were dried and eluted with 15 μL distilled H_2_O.

Library was constructed according to the manufacturer’s protocol (TD502, Vazyme). Briefly, 5 ng DNA obtained from earlier reaction was added to TTBL and TTE Mix V5 and incubated at 55℃ for 10 min. Once the reaction was completed, 5 μL TS buffer was added immediately, and the mixture was incubated for 5 min at room temperature. DNA fragments were amplified for 10-13 cycles to generate the library. Size selection was performed by adding 31 μL of DNA clean beads into 50 μL of PCR product and incubated for 5 min at room temperature. The supernatant was added into a new tube, 7.5 μL of DNA clean beads was added and followed by incubation at room temperature for 5 min. The beads were washed, dried and eluted with 20 μL distilled H_2_O, followed by sequencing as 150-bp paired-end reads on a platform Novaseq.

### scRNA-seq

Minced hippocampal tissues (∼2-4 mm^3^) were digested for 30min at 37℃ in the Papasin 20 U/mL enzymatic solution. The single-cell suspensions were filtered through a 70-μm cell strainer (Miltenyi Biotech) and then treated with the Red Blood Cell Lysis Solution (Miltenyi Biotech). The viability of cells was determined by using Countstar Rigel (Alit Biotech), and dead cell removal was carried out depending on the viability using the Dead Cell Removal Kit (Miltenyi Biotech). Finally, the cells were resuspended in 1x PBS (Invitrogen) supplemented with 0.04% BSA at a final concentration of 700-1200 cells/μL and were processed with the 10x Chromium Single Cell 3’ Kit (v3.1 PN:1000268) as per the manufacturer’s instructions. The cell suspension was loaded into 10x Chromium Chip (v3.1 PN:1000120) and barcoded with a 10x Chromium X. RNA from the barcoded cells was subsequently reverse-transcribed, amplified and prepared into sequencing libraries with 10x Library Construction Kit (v3.1 PN:1000190 or v4 PN:1000694) according to the manufacturer’s instructions. Sequencing was performed with Illumina NovaSeq with 150-bp paired-end reads at Novogene Bioinformatics Technology.

### FACS purification of mitotic DG NSCs

FACS purification of mitotic DG NSCs was carried out by adapting a newly developed BISMIB pipeline^15,84^. Briefly, cell suspension was briefly fixed with 0.1% formaldehyde for 10 min at room temperature and quenched with 0.125 M glycine for 5 min. The cells were then incubated with mouse anti-pH3-488 antibody (primary antibody conjugated with Alexa Fluor® 488) (1:200, Abcam, ab197502) in antibody buffer (Vazyme) at 4℃ for 2.5 h. After washing with cold PBS once, the cell suspension was filtering with 30 µm mesh before being analyzed by the Aria III based on pH3 immunofluorescence and cellular sizes. Mitotic and interphase NSCs were sorted based on the presence or absence of pH3-488 immunofluorescence respectively in wash buffer (Vazyme).

### CUT&Tag-seq

The CUT&Tag assays were carried out using Hyperactive Universal CUT&Tag Assay Kit for Illumina (TD903 and TD904, Vazyme). In brief, the NSCs and neural progenitors (same number of cells were used for control and experimental groups) were washed and re-suspended in wash buffer, before being incubated with ConA beads at room temperature for 10 min. The beads were incubated with primary antibody (1μg) in antibody buffer at 4℃ overnight, followed by incubation with secondary antibody (1:100) in Dig-wash buffer at room temperature for 1 h. After that, hyperactive pA/G-transposon containing buffer was added and incubated at room temperature for 1 h. TTBL buffer was added to activate transposase and incubated at 37℃ in thermomixer for 60 min. Protease K, buffer L/B and DNA Extract beads were then added to the sample, vortexed and incubated at 55℃ for 10 min. After washes, beads were dried. DNA fragments were eluted with 20 µL H_2_O.

Library amplification and product purification were performed using TruePrep Index Kit V2 for Illumina (TD202 and TD203, Vazyme) with the standard protocol. Size selection was then performed using VAHTS DNA Clean Beads (N411, Vazyme), in which 100 µL of DNA clean beads was added to 50 µL of PCR product, incubated for 10 min at room temperature. After washes, libraries were sequenced as 150-bp paired-end reads on a platform Novaseq.

The primary antibodies used for CUT&Tag analysis were rabbit anti-Prox1, rabbit anti-Fezf2, mouse anti-H3K27me3 (Abcam), rabbit anti-H3K4me3 (Abcam), rabbit anti-SUZ12; rabbit anti-EZH2 and rabbit anti-IgG (Abcam). For mitotic CUT&Tag, since the mouse anti-pH3 antibody used for FACS is a primary antibody conjugated with Alexa Fluor dye, CUT&Tag proceeds without the addition of a secondary antibody.

### Fluorescence recovery after photobleaching (FRAP) analysis

FRAP was performed using the Leica SP8 system with laser at 488 nm. Laser power for bleaching was attenuated to 40% using the microscope’s bleaching point mode. Images were collected every 10 s for 200 s after photobleaching. Note that some data that were obviously deviated due to changes of focal plane were not included in the statistics.

### Coimmunoprecipitation

48 h after transfection, HEK293T cells were harvested, washed and resuspended in lysis buffer [50 mM Tris-HCl (pH 8.0); 120 mM NaCl; 5 mM EDTA; 1% NP-40; 10% glycerol; protease inhibitor cocktail (Sigma-Aldrich); 2 mM Na_3_VO_4_] and kept on ice for 20 min. Cell extracts were sonicated with Bioruptor Plus (Biosense) at 4°C, clarified by centrifugation, and proteins immobilized by binding to anti-FLAG M2 (Sigma-Aldrich, A2220) affinity gel for 4 h at 4°C. Beads were washed and proteins recovered directly in SDS-PAGE sample buffer. Rabbit anti-FLAG (Cell Signaling Technology, 2368), rabbit anti-HA (Cell Signaling Technology, 3724) and Rb anti-GFP (Abcam, ab290) were used for western blot analysis.

### Quantification and statistical analysis

#### RNA-seq analyses

Raw FASTQ reads were trimmed using Trim Galore (v0.6.10, with parameters --paired --quality 20 --stringency 3 -j 4) to eliminate sequencing adaptors and low-quality bases. Quality of reads before and after trimming were monitored using FastQC (v0.11.9) and MultiQC (v1.14). Trimmed reads were then aligned to mouse genome (Version: mm10) using HISAT2 (v2.2.1) and processed with Samtools (v1.6) to generate bam files. FeatureCounts (v2.0.3) were used to generate raw count matrix for ncbiRefSeq genes. Further analyses were performed in R(v4.2.1). Differentially expressed genes (DEGs) were called using DESeq2 (v1.38.3). Log2 TPM (transcripts per kilobase million) normalization was used for heatmap visualization with ComplexHeatmap (v2.14.0). GO analysis was performed using clusterProfiler (v4.8.2), org.Mm.eg.db (v3.16.0) and ggplot2 (v3.4.2).

#### scRNA-seq analyses

Cellranger (v7.1.0) was used for alignment, filtering and UMI counting of single-cell FASTQs. The reads were mapped to the mm10 reference genome (refdata-gex-mm10-2020-A from Cellranger). Scrublet (v0.2.3) was used to detect and remove potential doublets. Filtered data were loaded into R (v4.2.1) for downstream analyses with Seurat (v4.3.0).

For E15.5 scRNA-seq data (Supplementary Fig. 2f–i), cells with gene numbers < 500 or gene numbers > 6000 or mitochondrial gene percentage > 15% were filtered out. SCTransform(variable.features.n = 3000) was used to normalize UMI count, and was followed by RunPCA(npcs = 50), FindNeighbors(dims = 1:30), FindClusters(resolution = 0.2), RunUMAP(dims = 1:30) to identify cell clusters. The 11 clusters of cells were annotated as 10 cell types based on expression of marker genes: NSC, progenitor, CA neuron, DG neuron, Cajal-Retzius cell, interneuron, microglia, oligodendrocyte precursor cells (OPC), endothelial cell and others. RNA velocity was calculated with velocyto (v0.17.17, run10x mode) and scVelo (v0.2.5).

For E15.5 control and *Prox1.6m^cKI^* scRNA-seq data (Fig. 5n), cells with gene numbers < 200 or gene numbers > 5000 or mitochondrial gene percentage > 5% were filtered out. The two samples were then merged and integrated with SCTransform(variable.features.n = 3000), RunPCA(npcs = 50), RunHarmony() from Harmony (v1.2.0). FindNeighbors(reduction = “harmony”, dims = 1:30), FindClusters(resolution = 0.2), RunUMAP(reduction = “harmony”, dims = 1:30) were used to identify cell clusters. The 12 clusters of cells were annotated as 11 cell types based on expression of marker genes: NSC, neural progenitor, CA neuron, DG neuron, Cajal-Retzius cell, interneuron, microglia, astrocyte progenitor, endothelial cell, mural cell and others.

### CUT&Tag analyses

Raw FASTQ reads were trimmed using Trim Galore (v0.6.10, with parameters --paired --quality 20 --stringency 3 -j 4) to eliminate sequencing adaptors and low-quality bases. Quality of reads before and after trimming were monitored using FastQC (v0.11.9) and MultiQC (v1.14). Trimmed reads were aligned to mouse genome (version: mm10) using Bowtie2 (v2.5.1) --very-sensitive--local mode. Picard tools (SortSam and MarkDuplicates) were used to mark and remove duplicate reads. Reads unmapped or with a mapping quality score < 10 were further filtered using Samtools (v1.6). For RPKM normalization, filtered bam files were transformed into bigWig or bedGraph files using deeptools bamCoverage (v3.5.1, with parameters --normalizeUsing RPKM --extendReads).

For spike-in normalization, trimmed reads were aligned to Escherichia coli genome (GenBank version: U00096.3) using Bowtie2 (v2.5.1) --very-sensitive--local mode. The scaling factor S was calculated as C (a constant) / E (fragments mapped to E. coli genome). Spike-in bigWig or bedGraph files were then generated using deeptools bamCoverage (v3.5.1, with parameters --scaleFactor scaling_factor S --normalizeUsing None --extendReads).

Heatmaps were generated with deepTools (v3.5.1) computeMatrix and plotHeatmap.

Prox1 and Fezf2 asynchronous binding peaks were called with Genrich (v0.6.1, -j -q 0.05). For mitotic Prox1 binding peaks, retained peaks were identified using Macs2 (v2.2.7.1, q=0.1). H3K4me3 binding peaks were identified with Macs2 narrow mode. H3K27me3 binding peaks were identified with Macs2 broad mode.

Macs2 bdgdiff was used to identify increased Fezf2 binding sites upon *Prox1* conditional knockout (Supplementary Fig. 3e–g), decreased Fezf2 binding sites from E13.5 to E14.5 in DG (Supplementary Fig. 3h-i) as well as increased H3K27me3 deposition upon Prox1 overexpression (Supplementary Fig. 5b).

DiffBind (v3.4.11) was used to quantify and compare deposition levels of H3K27me3 and H3K4me3 on specific genes (Fig. 6b).

All peak regions were loaded in R (v4.2.1) for further analysis. ChIPseeker (v1.34.1), org.Mm.eg.db (v3.16.0), ggplot2 (v3.4.2), clusterProfiler (v4.8.2) were used for peak annotation and visualization.

### Statistical methods

Quantification of the mitotic retention index of Prox1, Fezf2, SUZ12, EZH2 and RBBP4 is modified from previous methods calculating the chromosome enrichment levels^15,85^. Briefly, brain slices are stained for pH3 to mark the chromosomes. Retention index is calculated as log_2_ ratio of the mean fluorescence intensity (F.I.) of a specific protein on the chromosomes over the mean fluorescence intensity of the protein outside of chromosomes:

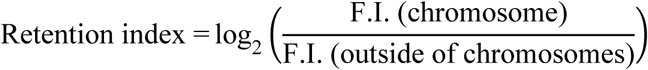

The retention index of positive values indicates enrichment on the chromosomes, whereas the retention index of negative values indicates exclusion from the chromosomes. Statistic data are presented as mean values ± SD (standard deviation).

## Data availability

The RNA-seq, CUT&Tag-seq and scRNA-seq data generated in this study have been deposited in the NCBI’s Gene Expression Omnibus (GEO) repository (GSE310910, GSE310650 and GSE310503). These data are publicly available as of the date of publication.

## Acknowledgements

We are grateful to Drs. Gordon Fishell, Fumio Matsuzaki, Chenglin Miao, Yulong Li, Yi Rao and Songhai Shi for mouse strains or reagents. We thank the National Center for Protein Sciences at Peking University, particularly Ms. Huan Yang for technical guidance on flow cytometry and Ms. Liqin Fu for technical support on microscope operation. We also thank Ms. Qiuju Zhu & Yonghong Song for general technical assistance, Ms. Yinuan Zha for plasmid construction, and members of the Song lab for discussions and help. This work was supported by the Ministry of Science and Technology of China (2022YFA1303100 to Y.S.), the National Natural Science Foundation of China (31771629 and 32371024 to Y.S.) and the Peking-Tsinghua Center for Life Sciences (to Y.S.). C.W. was supported in part by the Postdoctoral Fellowship of Peking-Tsinghua Center for Life Sciences.

## Author Contributions

C.W. and Y. Song conceived experiments. C.W., H.Y., H.L., X.L., T.L. and Z.C. performed mouse genetics, imaging, molecular biology, biochemical and multi-omics experiments. J.L. and J.C. performed bioinformatics analysis. Drs. Y. Shen and Y.Z. provided important technical guidance. C.W. and Y. Song wrote the manuscript with contributions from all authors.

## Additional information

Correspondence and requests for materials should be addressed to Yan Song.

---

**Supplementary Fig. 1.**
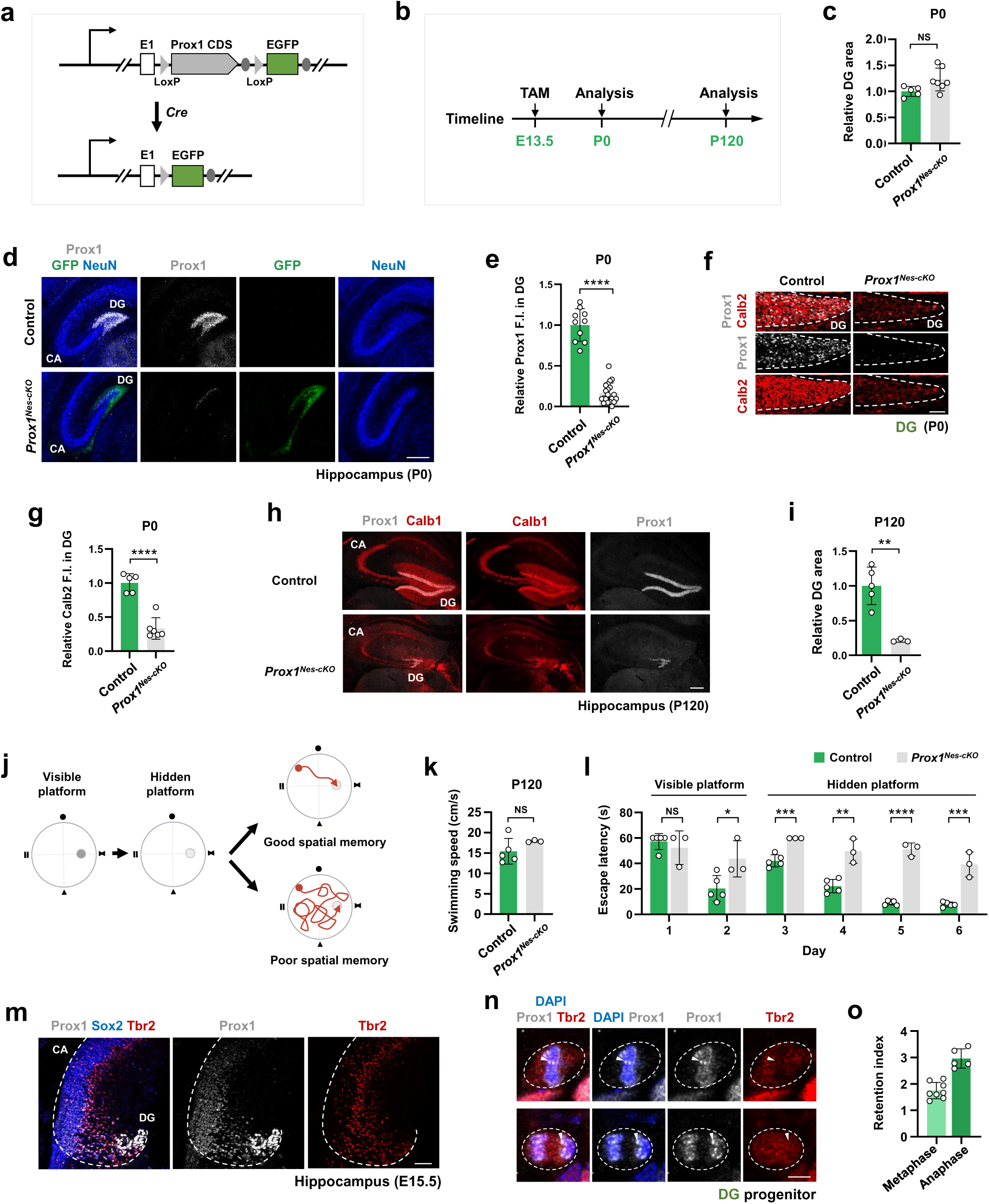
NSC lineage-specific knockout of Prox1 leads to severe DG malformation and spatial memory deficits. (**a**) A schematic diagram showing the genetic deletion strategy for generating *Prox1* conditional knockout mice. (**b**) A schematic diagram of the experimental design. TAM: tamoxifen injection. (**c**) Quantification of the relative DG area in control (*Prox1^fl/fl^*) and *Prox1^Nes-cKO^* hippocampus at P120. n=5 and 7 brain sections, respectively; NS, not significant; Student’s t-test. (**d–g**) Representative images showing control and *Prox1^Nes-cKO^* hippocampus at P0 stained for Prox1 and NeuN (**d**) or Prox1 and Calb2 (**f**), and quantification of relative Prox1 (**e**; n=10, 20 brain sections) and Calb2 (**g**; n=5, 6 brain sections) fluorescent intensity (F.I.). ****p < 0.0001; Student’s t-test. Scale bar, 200 μm (**d**); 50 μm (**f**). (**h**, **i**) Representative images showing control and *Prox1^Nes-cKO^* hippocampus at P120 stained for Prox1 and Calb1 (**h**), and quantification of relative Calb1 fluorescent intensity F.I. (**i**). n=5, 3 animals, respectively; **p < 0.01; Student’s t-test. Scale bar, 200 μm. (**j–l**) Morris water maze assay. A schematic diagram of the experimental design is shown in (**j**). Quantifications of the swimming speed (**k**) and escape latency (**l**) of control and *Prox1^Nes-cKO^* mice at P120. n=5, 3 animals, respectively; NS, not significant, *p < 0.05, **p < 0.01, ***p < 0.001, ****p < 0.0001; Student’s t-test. (**m**) Representative images showing hippocampus at E15.5 stained for Prox1, Sox2, and Tbr2. Scale bar, 50 μm. (**n**, **o**) Representative images showing the distribution pattern of Prox1, Tbr2 and DAPI in E15.5 DG progenitors at metaphase (up) and anaphase (bottom) (**n**) and quantifications of the retention index (**o**). Dashed lines indicate progenitor cell surface and arrowheads indicate Prox1 foci associated with the chromosomes. n=8, 5 cells, respectively. Scale bar, 5 μm.

**Supplementary Fig. 2.**
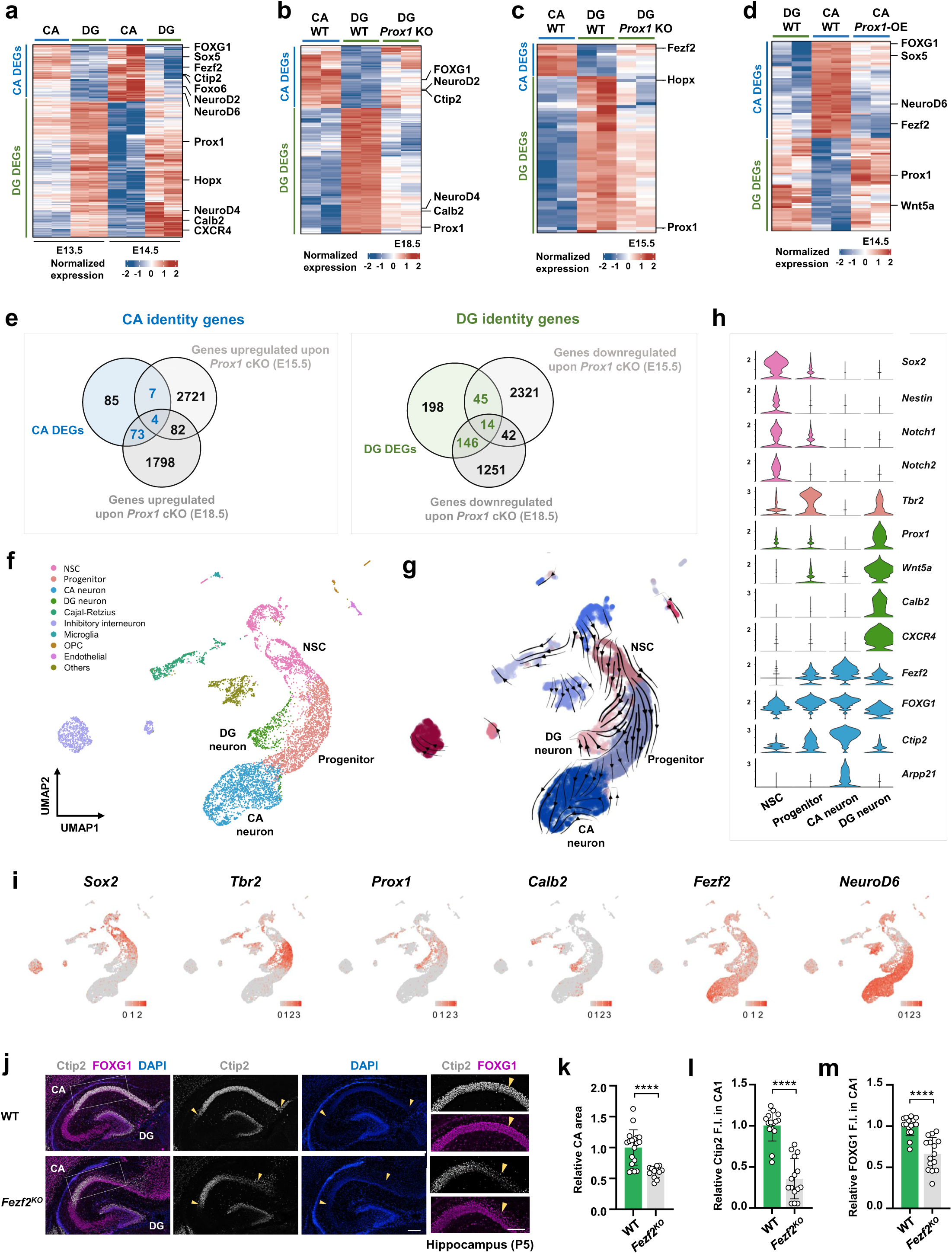
Prox1 defines DG lineage identity by suppressing the alternative CA identity. (**a**) Heat map illustrating the expression of CA and DG DEGs (differentially expressed genes) in CA and DG regions at E13.5 and E14.5. (**b**, **c**) Heat map showing the expression of CA and DG DEGs in WT CA, WT DG and *Prox1^Nes-cKO^* DG at E18.5 (**b**) and in WT CA, WT DG and *Prox1^Emx1-cKO^* DG at E15.5 (**c**). (**d**) Heat map illustrating the expression of CA and DG DEGs in WT DG, WT CA and CA overexpressing Prox1 (IUE: E13-E14.5). OE: overexpression. (**e**) Venn diagram showing the overlap between CA DEGs and genes upregulated upon *Prox1* cKO (left) and the overlap between DG DEGs and genes downregulated upon *Prox1* cKO (right). (**f–i**) scRNA-seq of early hippocampal development at E15.5. UMAP plot depicting annotated cell types (**f**). RNA velocity demonstrating streams of DG and CA neurogenesis (**g**). Violin plots (**h**) and feature plots (**i**) showing expression of cell fate and lineage identity genes in early hippocampal development. (**j–m**) Representative images of WT and *Fezf2^KO^* DG at P5 stained for Calb2, FOXG1 and DAPI (**j**) and quantifications of relative CA area (**k**), Ctip2 (**l**) and FOXG1 (**m**) fluorescent intensity in CA1 regions. n=19, 12 brain sections, respectively (**k**), n=15, 15 brain sections, respectively (**l, m**); ****p < 0.0001; Student’s t-test. Scale bar, 200 μm (left) and 100 μm (right).

**Supplementary Fig. 3.**
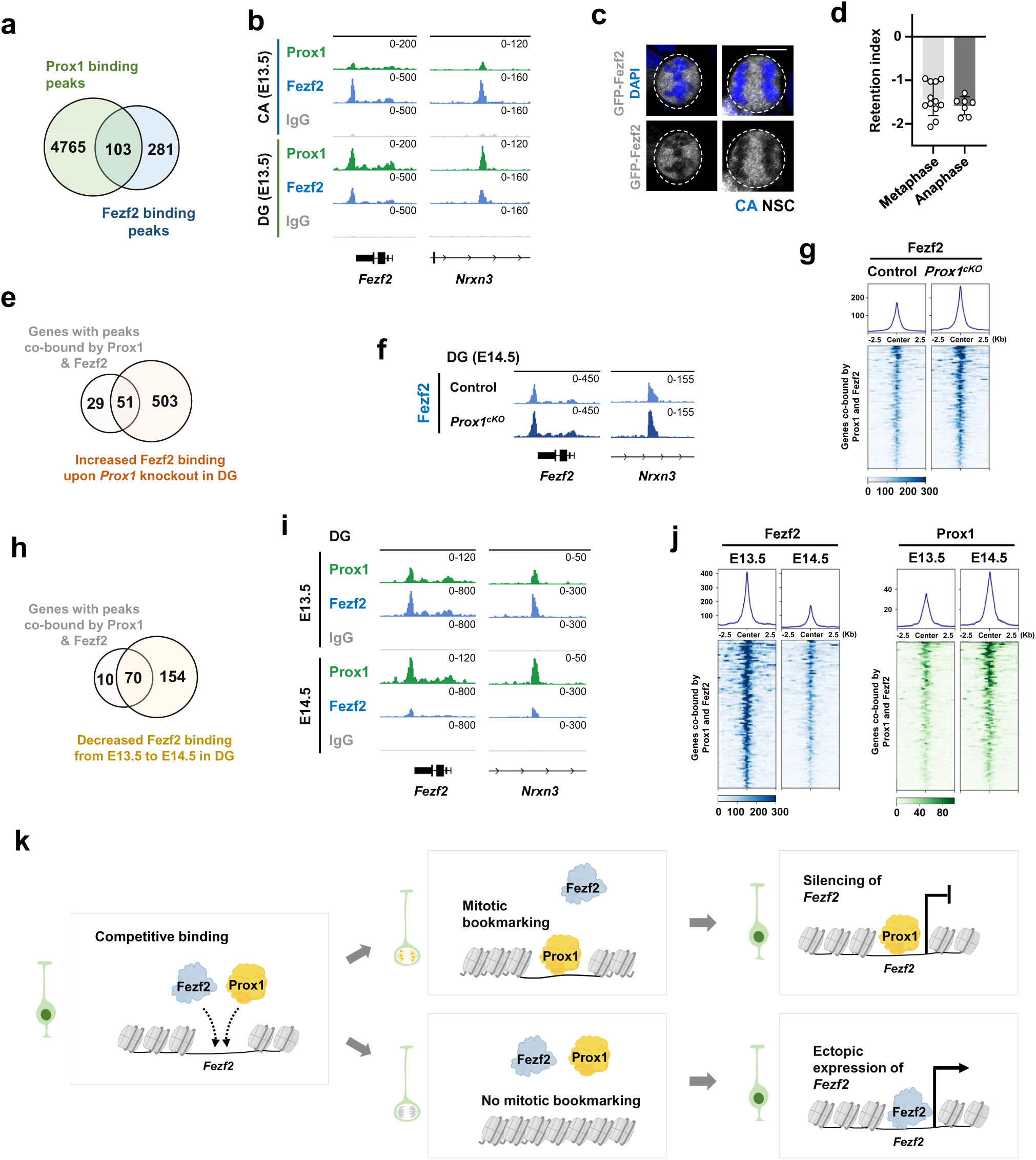
Mitotic retention provides Prox1 advantages in its competitive target gene binding with Fezf2. (**a**) Venn diagram showing the overlap of Prox1 and Fezf2 binding peaks in DG NSCs at E13.5. (**b**) Genome browser images of CUT&Tag for Prox1 and Fezf2 tracks at *Fezf2* and *Nrxn3* gene locus in CA (top) and DG (bottom) NSCs at E13.5. (**c**, **d**) Representative images showing the distribution pattern of GFP-Fezf2 and DAPI in CA NSCs at metaphase (left) and anaphase (right) (**c**) and quantifications of retention index (**d**). n=12, 7 cells. Scale bar, 5 μm. (**e**) Venn diagram showing the overlap of genes with peaks co-bound by Prox1 and Fezf2 and genes with increased Fezf2 binding affinity upon Prox1 knockout. (**f**) Genome browser images of CUT&Tag for Fezf2 tracks at *Fezf2* and *Nrxn3* gene locus in control and *Prox1^Emx1-cKO^* DG NSCs at E14.5. (**g**) Heat map showing Fezf2 binding peaks in control and *Prox1^Emx1-cKO^* DG NSCs. (**h**) Venn diagram showing the overlap of genes with peaks co-bound by Prox1 and Fezf2 and genes with decreased Fezf2 binding from E13.5 to E14.5. (**i**) Genome browser images of CUT&Tag for Prox1, Fezf2 and IgG tracks at *Fezf2* and *Nrxn3* gene locus in DG NSCs at E13.5 and E14.5. (**j**) Heat maps showing Fezf2 (left) and Prox1 (right) binding peaks in DG NSCs at E13.5 and E14.5. (**k**) A schematic diagram showing that mitotic retention of Prox1 allows it to outcompete Fezf2 in binding to the *Fezf2* gene locus in DG NSCs.

**Supplementary Fig. 4.**
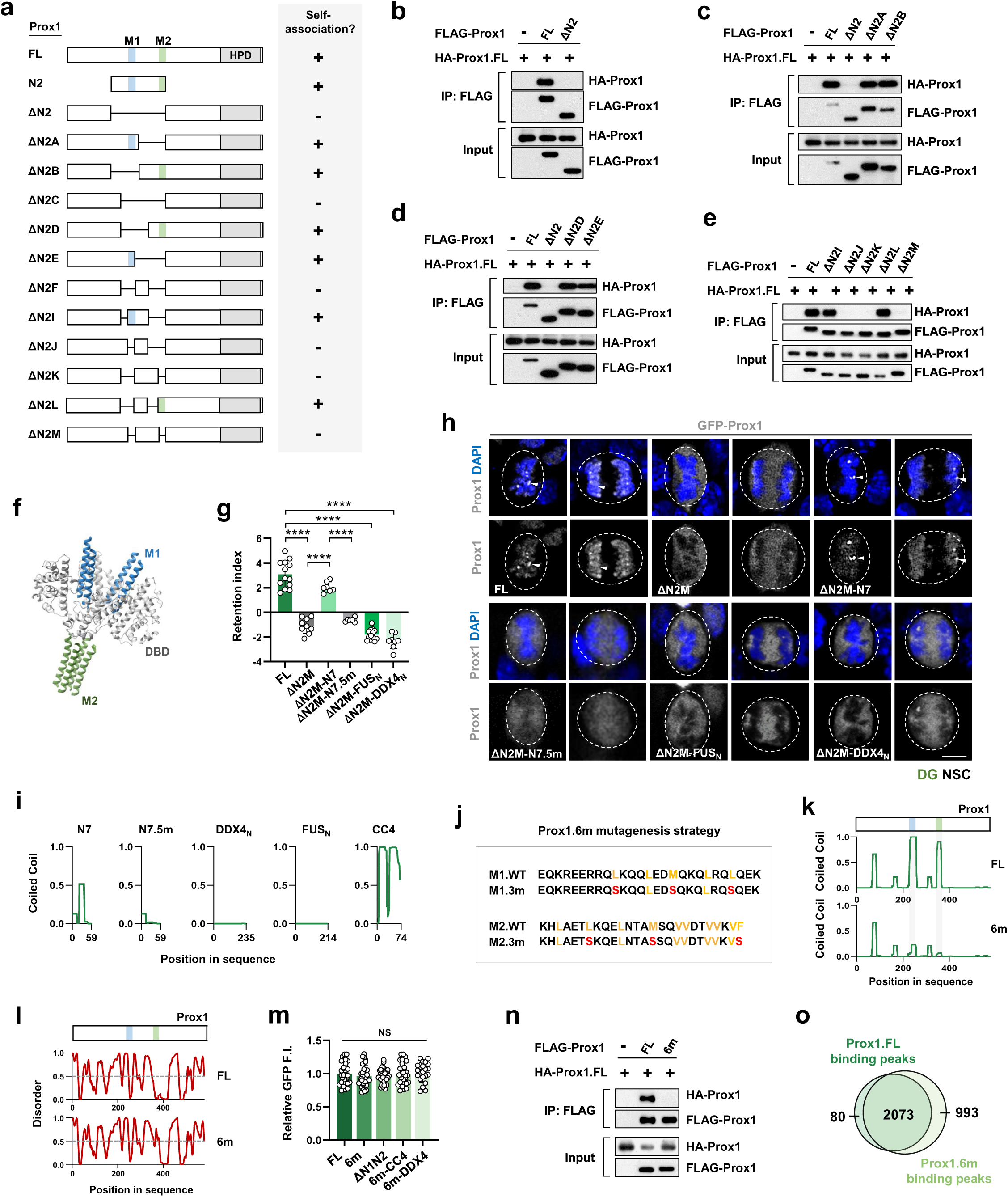
The 6m mutagenesis specifically impairs the mitotic retention ability of Prox1. (**a**) A schematic summary of the self-association ability of Prox1 variants. (**b–e**) Coimmunoprecipitation (coIP) of full length (FL) or truncated FLAG-tagged Prox1 with HA-tagged Prox1 in HEK293T cells. In these and subsequent panels, GFP served as a negative control and input represents 4% of total in coIP experiments. (**f**) Alphafold3 prediction of a Prox1 tetramer. The M1 and M2 motifs are in blue and green respectively. (**g**, **h**) Representative images showing the distribution pattern of GFP-Prox1 variants in DG NSCs at metaphase or anaphase (**h**) and quantifications (**g**). Dashed lines indicate NSC cell surface and arrowheads indicate Prox1 foci associated with the chromosomes. n=13, 9, 7, 6, 13, 8 cells, respectively; \**p* < 0.05, \*\*\*\**p* < 0.0001; Student’s t-test. Scale bar, 5 μm, (**i**) Coiled-coil prediction using PCOILS for protein domains used in restoration assay. (**j**) Mutagenesis strategy for Prox1.6m. (**k, l**) Coiled-coil prediction using PCOILS (**k**) and intrinsic disordered region (IDR) prediction using PONDR (**l**) for protein domains in Prox1.FL and Prox1.6m. (**m**) Quantifications of relative GFP-Prox1 fluorescent intensity (F.I.) in Fig. 3g. (**n**) CoIP of FLAG-tagged Prox1.FL or Prox1.6m with HA-tagged Prox1 in HEK293T cells. (**o**) Venn diagram showing the overlap of Prox1.FL and Prox.6m binding peaks.

**Supplementary Fig. 5.**
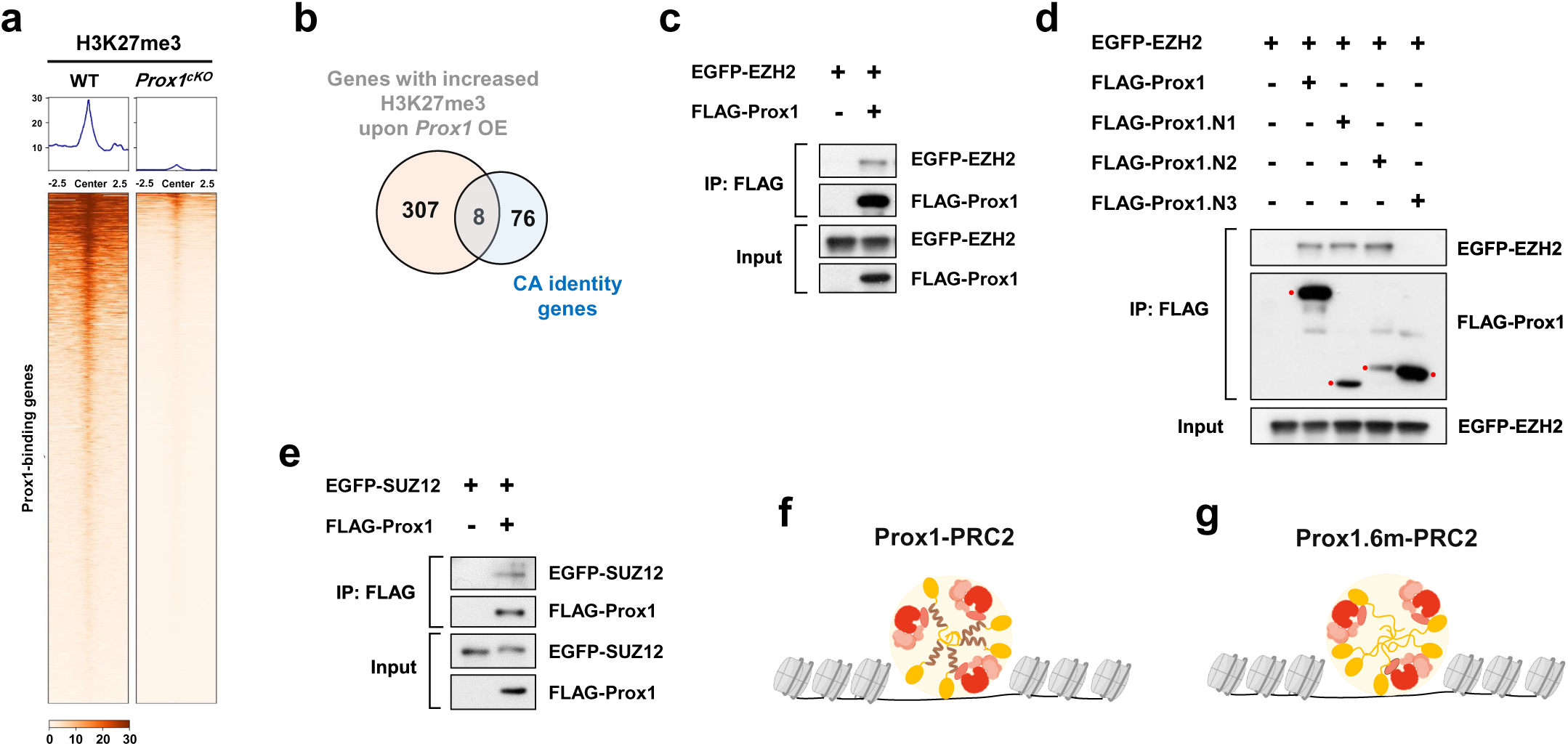
Prox1 forms protein complex with PRC2. (**a**) Heat map demonstrating Prox1-binding genes with significantly reduced H3K27me3 deposition upon *Prox1* knockout in DG lineages. (**b**) Venn diagram showing the CA identity genes with significantly increased H3K27me3 deposition upon Prox1 overexpression in CA lineages. (**c**, **d**) CoIP of full length or fragments of FLAG-tagged Prox1 with EGFP-tagged EZH2 in HEK293T cells. (**e**) CoIP of FLAG-Prox1 with EGFP-SUZ12 in HEK293T cells. (**f**, **g**) Schematic drawings of chromatin binding by Prox1-PRC2 (**f**) or Prox1.6m-PRC2 (**g**) co-condensates.

**Supplementary Fig. 6.**
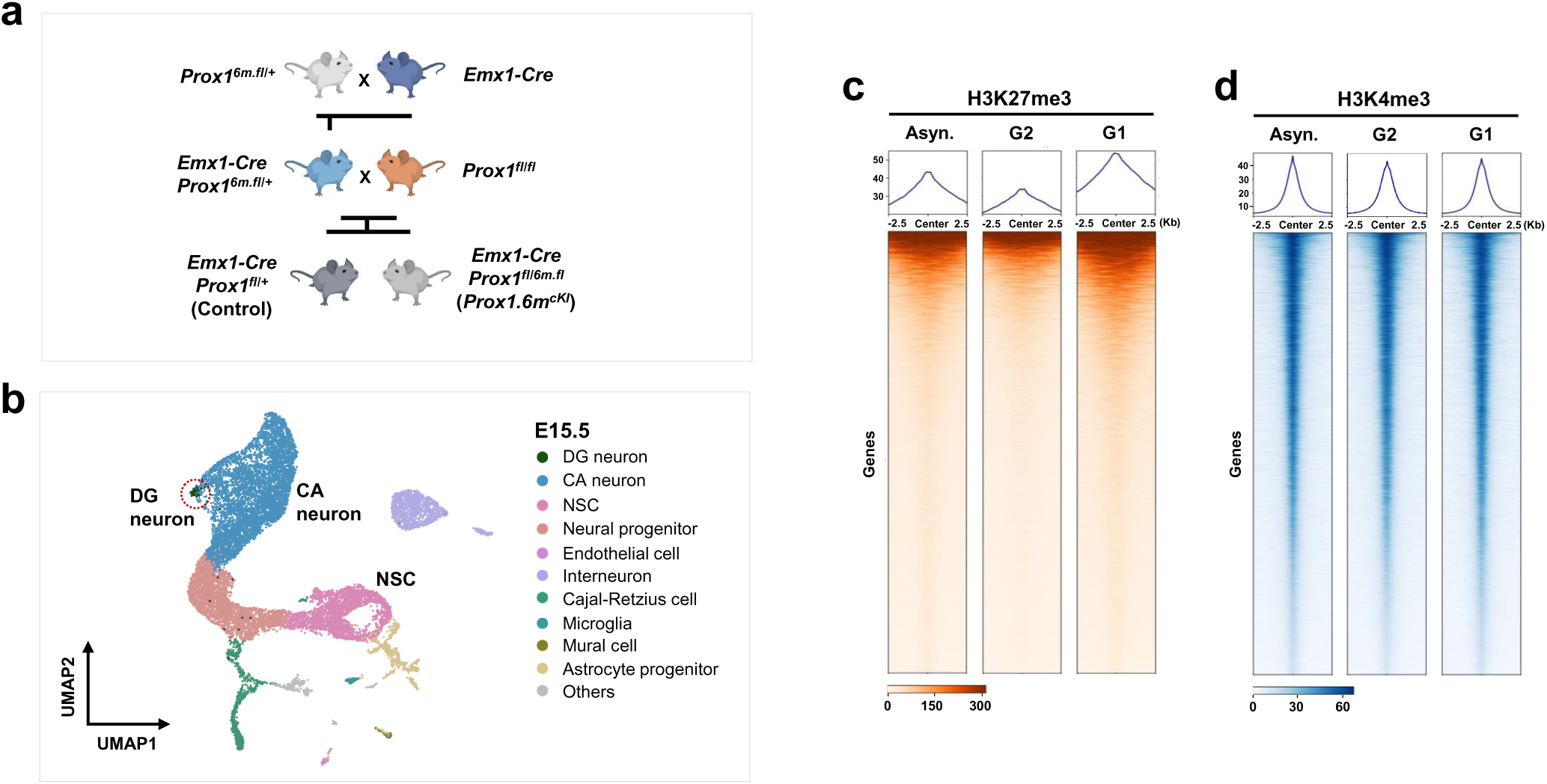
Functional analysis of Prox1 mitotic bookmarking and genome-wide histone bivalency in early hippocampal development. (**a**) A schematic diagram illustrating the generation of NSC lineage-specific *Prox1^6m^* knock-in mice (referred to as *Prox1.6m.^cKI^*) with NSC lineage-specific *Prox1* knock-out mice by *Emx1-Cre* mediated recombination. (**b**) UMAP plot showing the integrated scRNA-seq datasets from control and *Prox1.6m^cKI^* hippocampus at E15. Dashed circle indicates Prox1^+^Calb2^+^ DG neurons. (**c**, **d**) Heat maps of H3K27me3 (**c**) or H3K4me3 (**d**) deposition in asynchronous (Asyn.), G2 or G1 DG NSCs at E13.5.

